# Marine metabolomics: a method for the non-targeted measurement of metabolites in seawater by gas-chromatography mass spectrometry

**DOI:** 10.1101/528307

**Authors:** Emilia M Sogin, Erik Puskas, Nicole Dubilier, Manuel Liebeke

## Abstract

Microbial communities exchange molecules with their environment that play a major role in global biogeochemical cycles and climate. While extracellular metabolites are commonly measured in terrestrial and limnic ecosystems, the presence of salt in marine habitats has hampered non-targeted analyses of the ocean exo-metabolome. To overcome these limitations, we developed SeaMet, a gas chromatography-mass spectrometry (GC-MS) method that detects at minimum 107 metabolites down to nano-molar concentrations in less than one milliliter of seawater, and improves signal detection by 324 fold compared to standard methods for marine samples. To showcase the strengths of SeaMet, we used it to explore marine metabolomes *in vitro* and *in vivo*. For the former, we measured the production and consumption of metabolites during culture of a heterotrophic bacterium that is widespread in the North Sea. Our approach revealed successional uptake of amino acids, while sugars were not consumed, and highlight the power of exocellular metabolomics in providing insights into nutrient uptake and energy conservation in marine microorganisms. For *in vivo* analyses, we applied SeaMet to explore the *in situ* metabolome of coral reef and mangrove sediment porewaters. Despite the fact that these ecosystems occur in nutrient-poor waters, we uncovered high concentrations of many different sugars and fatty acids, compounds predicted to play a key role for the abundant and diverse microbial communities in coral-reef and mangrove sediments. Our data demonstrate that SeaMet advances marine metabolomics by enabling a non-targeted and quantitative analysis of marine metabolites, thus providing new insights into nutrient cycles in the oceans.

**Importance:** The non-targeted, hypothesis-free approach using metabolomics to analyzing metabolites that occur in the oceans is less developed than for terrestrial and limnic ecosystems. The central challenge in marine metabolomics is that salt prevents the comprehensive analysis of metabolites in seawater. Building on previous sample preparation methods for metabolomics, we developed SeaMet, which overcomes the limitations of salt on metabolite detection. Considering the oceans contain the largest organic carbon pool on Earth, describing the marine metabolome using non-targeted approaches is critical for understanding the drivers behind element cycles, biotic interactions, ecosystem function, and atmospheric CO_2_ storage. Our method complements both targeted marine metabolomic investigations as well as other ‘omics’ (e.g., genomics, transcriptomics and proteomics) level approaches by providing an avenue for studying the chemical interaction between marine microbes and their habitats.

## Introduction

Marine microorganisms produce and stabilize the largest pool of organic carbon on Earth by exchanging molecules with their environment (1, 2). Marine microbes are also the basis for maintaining the long term storage of carbon dioxide (CO_2_) in the oceans, which plays a complex role in biogeochemical cycles with uncertain implications for global climate (3). While metagenomic and metatranscriptomic studies of the ocean, driven by low sequencing costs and projects like Tara Oceans (4), have deepened our knowledge of the identity and activity of marine microbes, these studies are limited in their ability to determine the molecules that contribute to the chemical complexity of marine habitats. New approaches are needed to permit equivalent surveys of the extracellular metabolome, or exometabolome of the ocean.

Exometabolomics provides an opportunity to directly characterize the molecular interaction between microbes and their environment by profiling the types of molecules cellular organisms secrete (5). In terrestrial and limnic systems, these studies have advanced our understanding of microbial communities in soil organic matter cycling (6, 7), overflow metabolism of cultivable microorganisms (8, 9) and chemical ecology of the environment (10, 11). While intracellular metabolomic analyses of tissues from marine microbial cells to invertebrates is becoming increasingly more common (12-14), the defining characteristic of marine habitats - high salt concentration - limits exometabolomic analyses of the oceans to studies that require salt removal prior to metabolite extraction (10, 15, 16).

Our knowledge of the metabolite composition of ocean habitats is restricted to methods that require sample preparation techniques that alter their molecular composition, or targeted approaches that measure a defined group of metabolites (17-19). The most common environmental profiling strategies in marine ecosystems rely on solid phase extraction (SPE) techniques to remove salt prior to mass-spectrometry (MS) analyses (20, 21). These studies have demonstrated the role of microbial communites in producing recalcitrant dissolved organic matter (DOM) and provided insights into their role in long term carbon storage (22). However, the removal of salt from marine samples using SPE is accompanied by the co-removal of small polar compounds, which are the primary components of the liable organic matter pool (17). Consequently, SPE-based studies can only detect about 50% of the compounds that make up the DOM pool from the ocean, and fail to detect the majority of compounds involved in the central metabolism of cells. Furthermore, current DOM analytical approaches remain largely inaccessible for the majority of research institutions and projects. This is largely due to high instrumentation costs for high-resolution MS (coupled to liquid-chromatography or with direct-infusion), large sample volume requirements, and the relatively low-throughput in data acquisition.

Gas chromatography (GC)- MS analysis, on the other hand, is a high-throughput and widely available analytical method that allows for the detection of primary metabolites, small molecules that occur in central metabolic pathways across biological systems (23, 24). High reproducibility coupled to the widespread availability of annotation resources make GC-MS the “workhorse” of analytical chemistry facilities. GC-MS has allowed the identification of metabolites associated with human disease (25), detection of compounds that serve as environmental cues in foraging (26), description of metabolic fluxes within and between cells (27), and is used for environmental profiling of soils and microbial activity on land (6, 28). Despite its the power of detecting metabolites involved in central metabolism, exometabolomic studies using GC-MS from marine habitats are absent due to the inhibitory effects of salt on sample analysis.

The ocean metabolome remains largely undefined, despite a growing field of research exploring the molecular composition of DOM (1, 2, 20, 21). To more efficiently decipher ocean metabolism, cost-effective, high-throughput, and untargeted workflows that can readily identify and quantify molecules from high salinity environments are critical. Here, we present SeaMet, a marine metabolomics method that builds on previous GC-MS sample derivatization methods to enable metabolite detection in seawater. Using SeaMet, we demonstrate how our method can enhance our understanding of microbial metabolism in culture experiments and profiling of marine habitats.

## Results and Discussion

SeaMet modifies the well-established two-step derivatization procedure, which permits the detection of non-volatile primary metabolites using GC-MS, and involves methoximation followed by trimethylsilylation (29). Like other GC-MS sample preparation techniques (30, 31), SeaMet removes liquid through vacuum drying prior to derivatization - a process that results in a salt pellet when working with marine samples, which restricts MS analysis. Our preliminary tests suggested that water locked within the dried salt crystals hindered the chemical reactions needed for GC-MS (**Fig. S1**). Our method overcomes this limitation by first eliminating residual water within the salt crystals and then extracting metabolites into the derivatization reagents (**Fig. 1A**).

**Figure 1.**
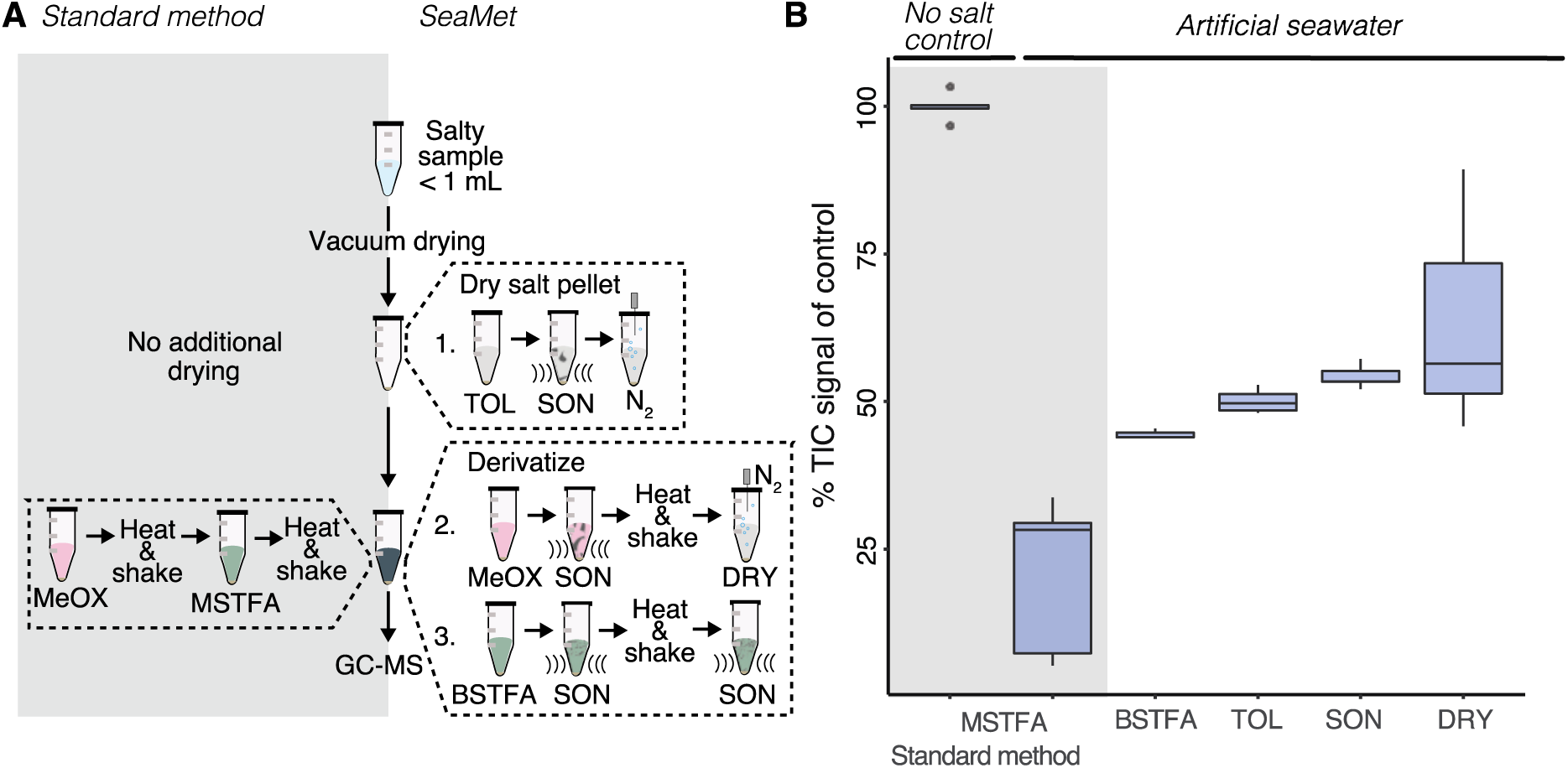
How SeaMet works. **A**, Modifications to the standard two-step methoximation (MeOX)-trimethylsilylation (TMS) derivatization protocol include key steps that enhance metabolite signal detection in seawater as shown in **B.** Steps modified from the standard method include a switch in derivatization reagents from MSTFA to BSTFA, further drying of the salt pellet using toluene (TOL) to remove water azeotropically, ultrasonication (SON) after the addition of TOL, MeOX, BSTFA, and after BSTFA derivatization, and drying (DRY) of the pyridine after the MeOX derivatization prior to BSTFA addition. **B**, Box plots showing changes in total ion chromatogram (TIC) signals after GC-MS data acquisition. Results are from a synthetic mixture of 45 metabolites representing a broad scope of metabolite classes (**Table S1**) dissolved in 0.5 mL of seawater (*n* = 5) relative to average of the no salt control.

We used a mixture of 45 different metabolites (**Table S1**) dissolved in artificial seawater to 0.4 mM to document the performance in metabolite detection of our method. Overall, SeaMet increased total signal intensity on average by 42% and up to 89% for high salinity samples in comparison to the standard GC-MS sample preparation (**Fig. 1B**). We first replaced the most commonly used trimethylsilylation reagent, N-methyl-N-(trimethylsilyl)trifluoroacetamide (MSTFA)(31), with one that is less susceptible to inhibition by water, N, O-Bistrifluoroacetamide (BSTFA), which resulted in higher metabolite signals (**Fig. S1B**). To eliminate water from the samples, we increased the speed-vacuum drying time from four to eight hours, and integrated a toluene drying step that is used in urine-based metabolomic analyses (30). We further enhanced metabolite signals by treating the salt pellet to a combination of ultrasonication and vortexing after the addition of toluene and both derivatization reagents, and following completion of the trimethylsilylation reaction. These steps break apart the salt crystals and release water into the toluene to enhance salt drying and metabolite extraction. Finally, following a recently described method for improving GC-MS metabolite detection regardless of sample type (32), we included an additional step between the methoximation and trimethylsilylation derivatization reactions and evaporated the first derivatization reagent under N_2_ gas (see **Fig. 1B** for total signal improvements of each step).

Overall, SeaMet allowed us to detect significant increases in metabolite abundances across molecular classes when compared to the standard method (adjusted *p*-value < 0.05; mean fold change across all ions = 323; **Fig. 2A; Fig. S2**). This included measurement of organic acids, amino acids, and fatty acids, as well as sugars (and their stereoisomers), sugar alcohols, and sterols (**Table S1**).

**Figure 2.**
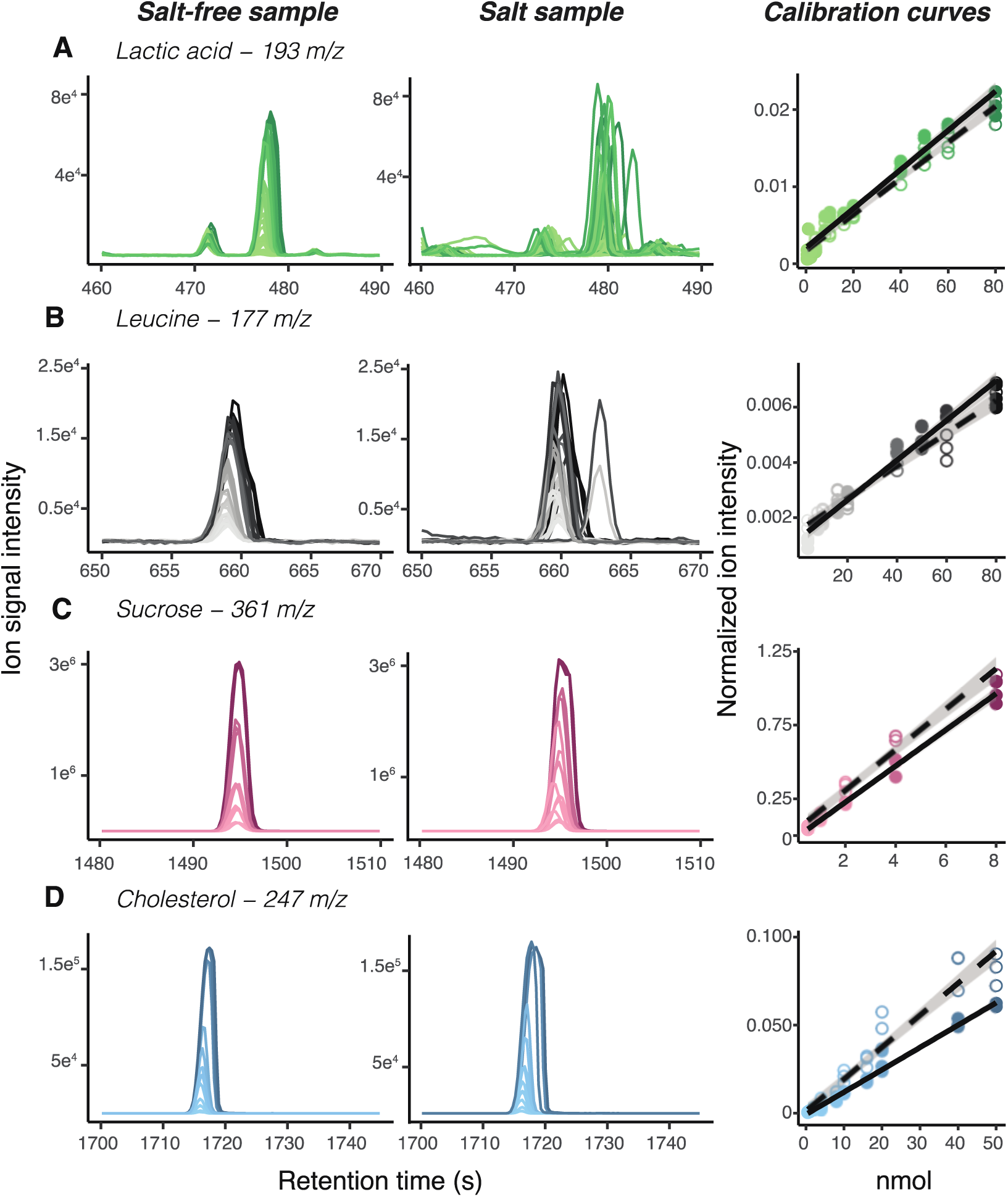
Metabolite detection and quantification in seawater using SeaMet in comparison to salt-free water. **A-D**, Extracted ion chromatograms for select metabolites in salt-free and artificial seawater demonstrate reproducible metabolite detection across concentration gradients, as shown in associated calibration curves on the right. Spectra and points are shaded in scale with sample concentrations, where more concentrated samples are represented by darker colors. Open circles = salt samples; filled circles = salt-free samples.

To determine the quantitative capabilities of SeaMet, we used a metabolite mixture (45 metabolites spanning 9 compound classes) and added different concentrations (from 0.0039 mM-to 0.4 mM) to seawater. Our detection limits were in the nano-molar range and comparable to those of targeted techniques for marine ecosystems that were developed to quantify single compounds from specific molecular classes (**Tables S2; Table S3**). In contrast to previously published techniques, which require at least an order of magnitude higher sample volumes, SeaMet only requires 0.5 mL to 1 mL of seawater for metabolite detection (17, 33). Using SeaMet, we measured 107 metabolite standards in seawater, representing major metabolite groups involved in primary metabolic pathways (**Table S4**). Our method provides reproducible quantification across metabolite classes (r^2^ > 0.7), and gives similar linearity and dynamic range in seawater samples compared to salt-free samples prepared with the standard GC-MS derivatization method (**Fig. 2A-H; Fig. S3)**. Moreover, we demonstrate that SeaMet reduces variation in ion detection for individual metabolites (Welches t-test *p*-value < 0.01 across all ions at 4 nmols; average % CV_salt_ = 20.2 ± 0.78, average % CV_salt-free_ =2 3.5 ± 0.72) compared to salt-free samples prepared with the standard GC-MS derivatization procedure (**Table S2**). The analytical characteristics of the 107 metabolites (**Table S4**) can be used for more sensitive, targeted GC-MS analyses or help in identifying metabolites in untargeted applications.

Given that SeaMet avoids SPE, we assessed how SPE sample treatment affects the ability to detect compounds in marine samples. We compared GC-MS profiles measured with SeaMet before and after salt removal using the most commonly used Bond Elut styrene-divinylbenzene (PPL) SPE-columns. Our analyses revealed that small polar compounds, such as sugars, sugar alcohols, amino acids, and organic acids, were co-removed with salt during SPE sample preparation (**Fig. 3B**). These results provide evidence that SeaMet captures compounds commonly missed by SPE-based exometabolomic approaches for marine samples. SeaMet thus expands the range of metabolites that can be measured by untargeted approaches beyond those currently used to characterize marine DOM, and contributes to advancing marine metabolomics.

**Figure 3.**
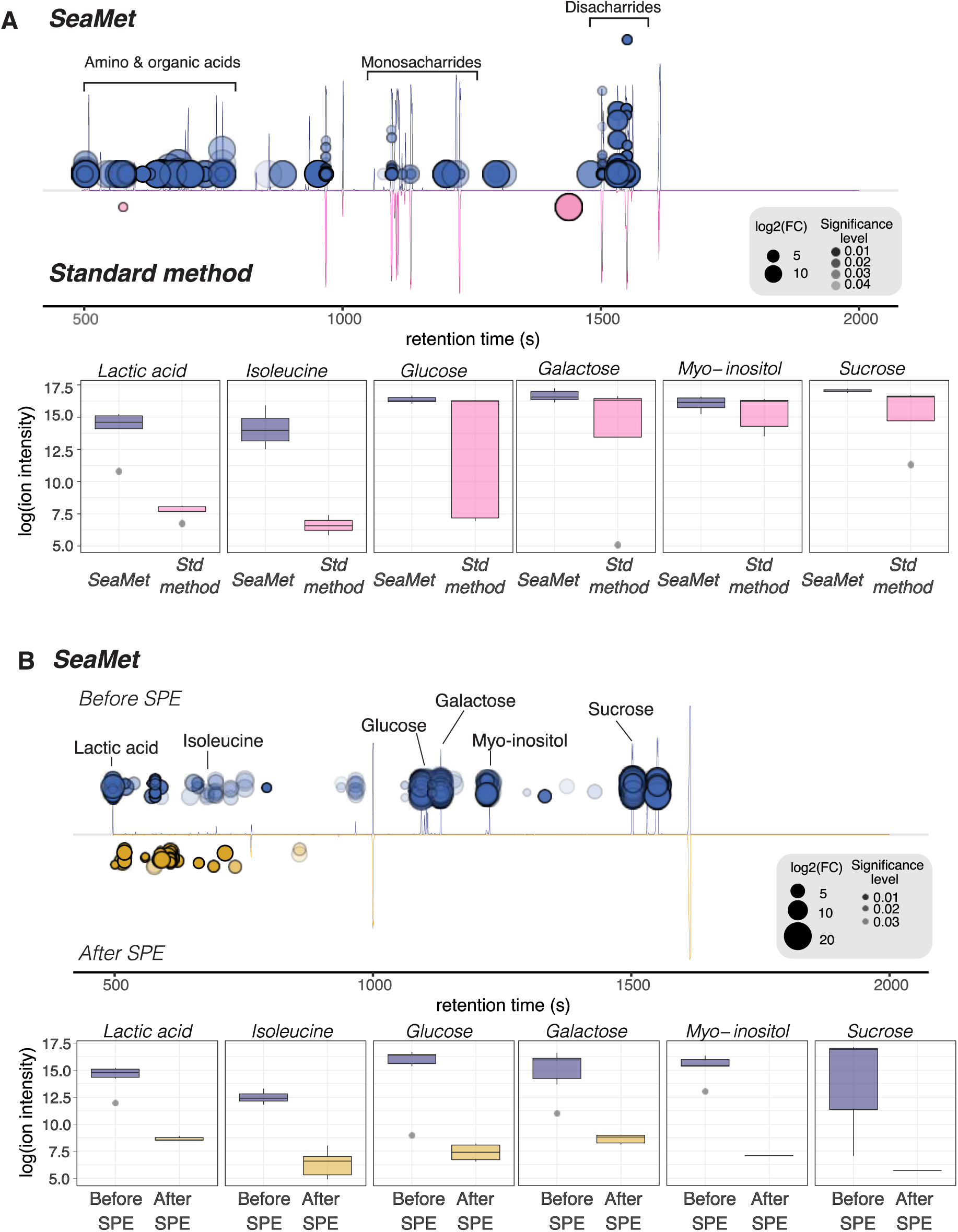
SeaMet enhances the detection of metabolites in marine samples. **A, B** Total ion chromatogram cloud plots from GC-MS profiles of metabolite mixtures indicate significant differences (Benjamini-Hochberg adjusted *p-value* < 0.05) between ion abundances when comparing **A**, SeaMet (blue; top) to the standard metabolite derivatization (pink; bottom) protocol for GC-MS samples and **B**, chromatograms using SeaMet on marine samples before (blue; top) and after (yellow; bottom) solid phase extraction (SPE). Individual compound box plots are also shown in **A** and **B** to highlight improvements in metabolite detection using SeaMet. For the cloud plots, larger bubbles indicate higher log2(fold changes) between groups and more intense colors represent lower t-test p-values when comparing individual feature (*m*/*z* ions) intensities. Samples prepared with SeaMet had high abundances of organic acids (lactic acid, succinic acid, and fumarate), amino acids (isoleucine, leucine, threonine and valine), sugar alcohols (myo-inositol and mannitol), and sugars (fructose, glucose, cellobiose, maltose, ribose, galactose, and sucrose) in comparison to SPE-based sample preparation. Representatives of each class are indicated in **B**. To show signal improvement using SeaMet, samples for both comparisons included authentic metabolite standards representing multiple chemical classes.

To demonstrate the power of SeaMet in characterizing the metabolism of marine bacteria, we monitored changes in the extracellular metabolome during growth of a heterotrophic Gammaproteobacterium, *Marinobacter adhaerens* that occurs in aggregation with diatoms throughout the North Sea. Using SeaMet, we simultaneously observed hundreds of metabolites and detected significant changes in the metabolite composition of marine culture medium during the bacteria’s initial growth phase (adjusted *p*-value < 0.05; **Fig. 4; Fig. S3).** The bacteria took up different carbon and nitrogen resources in a cascade-like fashion, and later in growth, began excretion of an undescribed compound (**Fig. 4C, D; Fig. S3**). By measuring multiple metabolite classes in a single analytical run, our results revealed that *M. adhaerens* preferentially took up amino acids over readily available sugar compounds (e.g., trehalose, **Fig. S3**). Previous proteomic results indicated that *M. adhaerens* had a high number of expressed amino acid uptake transporters (34). Our results expand on these findings by i) highlighting which amino acids the *M. adhaerens* prefers, ii) providing experimental evidence that this heterotroph does not take up sugars, despite the genomic ability to use them in their metabolism (35), and iii), showcasing that *M. adhaerens* participates in the successional uptake of resources. Successional dynamics in substrate use is a common energy conservation mechanism in bacteria (36) and affects central carbon and nitrogen dynamics during growth. *M. adhaerens*, like many other bacteria, participates in the release of organic carbon, which can be metabolized by other microorganisms or will contribute to the complexity of refractory DOM.

**Figure 4.**
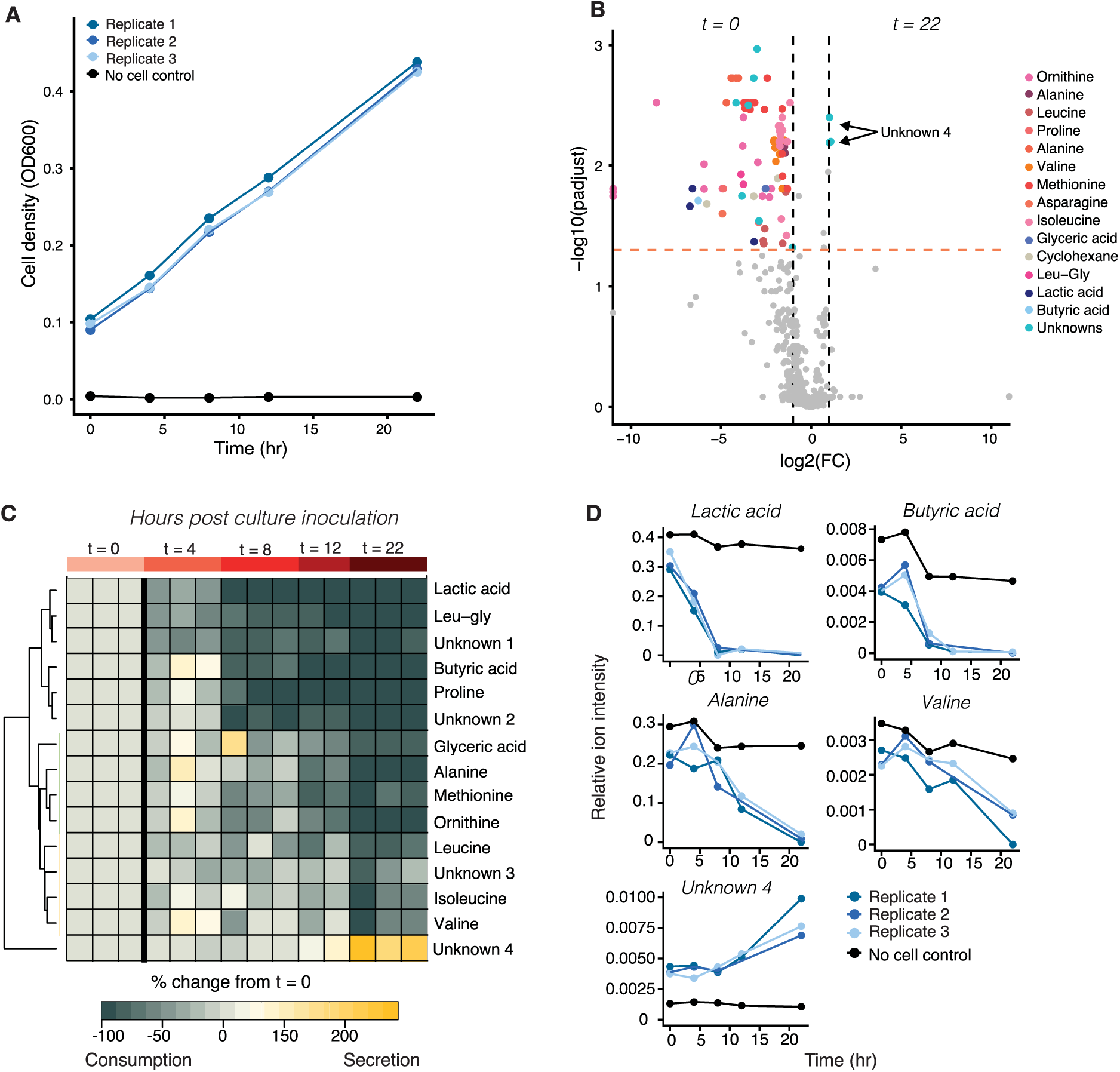
Metabolite consumption and excretion during culture of the marine heterotroph. *Marinobacter adhaerens*. **A**, Cell densities increased during the first 22 hours of culture growth in Marine Broth. **B**, Volcano plot showing differences in ion abundances in cell growth media between the initial and final (22 hour) sampling time points. Variables exhibiting high fold change values (log2(fold change) > 2) and significant differences (*p*-adjusted < 0.05) between the two sampling time points are colored according to their metabolite database (NIST) annotation. **C**, A heatmap of metabolite abundances after 22 h relative to starting conditions indicates some compounds, like the dipeptide leucine-glycine (leu-gly), and lactic acid were taken up before others, such as branch chain amino acids. After 12–22 hours of growth, the bacteria excreted an unknown compound (Unknown 4). Hierarchical clustering shows groups of metabolites that changed significantly during growth (left hand colored bars, B.H. adjusted p-value < 0.05; fold change > 2). These metabolite groups represent successive stages in *M. adhaerens* consumption and production of marine broth components. **D**, Relative ion abundances over time for select metabolites from each cluster group shown in **C.** The blue lines represent biological replicate cultures while the black line shows results from a control sample with no cell addition. Low variation among biological replicates highlights the reproducibility of SeaMet.

Given that other exometabolomic methods for marine samples either miss major compound classes due to sample pre-treatment (e.g., SPE based sample preparation) or are targeted approaches that can only measure a few metabolite groups in a given run, it is likely these observations in *M. adhaerens* physiology would have been obscured. Give the ease in applying our method to culture studies, it is possible to integrate SeaMet with other “omics” approaches to help illuminate microbial physiology in the marine environment. By identifying and quantifying metabolites that are consumed and excreted in cultivable marine bacteria, our method expands our understanding of key primary compounds involved in the transformation of organic matter in the ocean.

To test the ability of our workflow to assess complex environmental metabolomes, we applied SeaMet to porewater samples from coralline and mangrove sediments. Coral reefs and mangroves, two globally important coastal ecosystems, contain many biological compounds that remain undescribed. It is essential to characterize the metabolome of these habitats to understand the role of these ecosystems in biogeochemical cycling.

Our approach detected 295 and 428 metabolite peaks from coralline and mangrove porewater profiles (**Fig. 5**), including sugars, amino acids, organic acids, fatty acids, and background signals. Diverse and abundant sugars from sediment porewaters adjacent to corals, as well as fatty acids from porewaters next to mangroves drove the observed significant differences between habitats (ADONIS *p*-value < 0.001, *R*^2^ = 0.514; **Fig. 5 and Table S5**).

**Figure 5.**
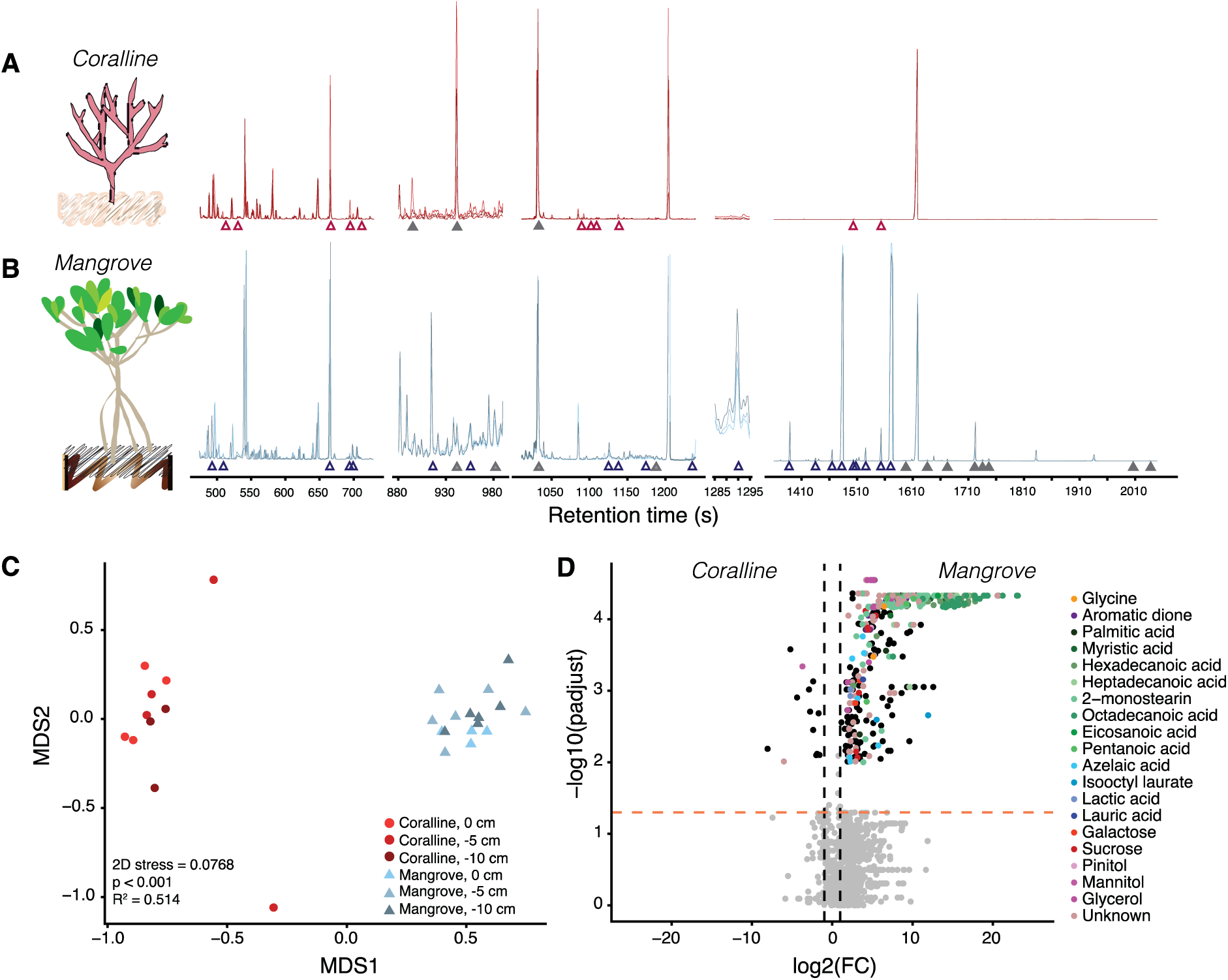
Metabolite profiles from marine habitats acquired with SeaMet. GC-MS metabolomic profiles from **A**, coralline and **B**, mangrove sediment porewaters showed high concentrations of identified metabolites (open triangles), e.g. fatty acids and sugars that explain multivariate differences in composition in **C**. Profiles also revealed unknown peaks (filled triangles) for which no matches were found in public databases (**Table S5**). **C**, Bray-Curtis informed non-metric multidimensional scaling analysis of sediment porewater metabolomic profiles from coralline (red) and mangrove (blue) habitats across sediment depths. ADONIS p-value and R^2^ showed a significant correlation between sampling location and metabolite composition. **D**, Volcano plot showing differences in ion abundances between habitats. Significant ions (*p*-adjusted < 0.05) with log2-fold change > 2 are shaded according to their metabolite database (NIST) annotation.

Given that corals and mangroves thrive in oligotrophic waters and their associated sediments harbor diverse, abundant and metabolically active microorganisms (37, 38), we were surprised to measure high levels of metabolites that are typically consumed in primary metabolism. Metabolomic analyses of marine sediments (in bulk) have also detected high abundances of primary metabolites (39, 40), suggesting sediment habitats – which are globally home to an estimated 2.9 X 10^29^ microbial cells (41) – contain many different types of metabolites that drive microbial community metabolism. These data call for a reexamination of carbon sequestration in coastal sediments using techniques that can identify and quantify the accumulation of liable metabolites.

Due to the technical difficulties of detecting metabolites in seawater, a large portion of ocean chemistry remains unannotated, reflecting one of the central challenges in metabolomics research (42). By providing a new method to measure a broad scope of the marine metabolome, we offer an avenue to identify molecules from marine environments and expand existing mass spectrometry databases that aim to characterize chemical space across ecosystems. As an example, our samples from sediment porewaters of mangroves and coral reefs revealed 11 metabolites driving variation between habitats that did not match public database entries (Fig. S6 and **Table S5**) (43, 44).

## Conclusions

SeaMet is a marine metabolomics workflow that enables the analysis of primary metabolism in the oceans. It is time efficient, allows the detection of diverse metabolite classes in a single run, and expands the analytical window for molecules that can be detected within marine samples. This advance enables untargeted metabolomics for marine ecosystems using a low-cost, easy to use GC-MS platform. Moreover, SeaMet is independent of MS instrumentation, allowing it to be combined with time-of-flight or Orbitrap MS detectors to provide faster analysis time and higher mass resolving power to improve metabolite identification. We expect our marine metabolomics workflow will enable the exploratory analysis of metabolites occurring in seawater and thereby advance our understanding of the ocean’s vast and largely unexplored metabolome.

## Materials and Methods

### Data availability

All metabolite profile data will be made publicly available at Metabolights (https://www.ebi.ac.uk/metabolights/) under identification numbers MTBLS826, MTBLS839, MTBLS843, MTBLS844, MTBLS848, and MTBLS849 (currently IN REVIEW) or by contact with the authors. Reviewer links: https://www.ebi.ac.uk/metabolights/reviewer5eb6b480436b019d9f1351a828ee7c3d

https://www.ebi.ac.uk/metabolights/reviewerd923ea1c3a53d000b97ccf383991032d

https://www.ebi.ac.uk/metabolights/reviewera9be9cf4-9a7d-4fff-98d5-c3d574c3b7f5

https://www.ebi.ac.uk/metabolights/reviewer08ce2d89-3945-45be-8a9b-4ea872fc86bf

https://www.ebi.ac.uk/metabolights/reviewer07a4ce73-1e8e-46aa-80c8-0c2f26411174

https://www.ebi.ac.uk/metabolights/reviewer2878413e6f8a6a883b27bdee8c1bbba6

### Reagents and experimental sample preparation

The derivatization chemicals, trimethylsilyl-N-methyl trifluoroacetamide (MSTFA) and N,O-Bis(trimethylsilyl)trifluoroacetamide (BSTFA) were obtained from CS-Chromatographie Service and pyridine from Sigma-Aldrich at >99.98% purity. Methoxyamine hydrochloride (MeOX; Sigma-Aldrich) aliquots were further dried at 60 °C in a drying oven for 1 h to remove residual moisture. Artificial seawater (ASW) was prepared within the range of natural salinity (36‰) by dissolving (per L of water) 26.37 g sodium chloride, 6.8 g magnesium sulfate heptahydrate, 5.67 g magnesium chloride hexahydrate, 1.47 g calcium chloride, 0.6 g potassium chloride, and 0.09 g potassium bromide. Following autoclave sterilization, pH was adjusted to 7.7 using sodium hydroxide. 1 mL of the following supplements and solutions were added: 150 mM monopotassium phosphate, 500 mM ammonium chloride pH 7.5, trace element solution, selenite-tungstate solution, vitamin solution, thiamine solution, B12 solution and 0.21 g sodium bicarbonate (45). Ultra-pure water (MQ) was prepared by purifying deionized water with an Astacus membraPure system (Astacus membraPure, 18.3 m*Ω* × cm 25 °C).

Metabolite standards were obtained from commercial sources (**Table S4**) and combined into mixtures in which each compound had a final concentration of 0.4 mM. Metabolite mixtures were prepared to (a) test the effect of salt and water on metabolite detection, (b) develop SeaMet, our marine metabolomics workflow, (c) compare metabolite detection before and after solid phase extraction (SPE) based sample preparation, and (d) to quantify the detection limits of specific compound classes (**Table S6**). Finally, multiple mixtures were prepared to document the retention times of 107 standards dissolved in ASW using SeaMet (**Table S4**). Sample aliquots for the above mentioned experiments were prepared by drying down 200 µL of the mixture in a speed vacuum concentrator (Eppendorf Concentrator Plus^(R)^, 2.5 h, 45°C, V-AQ) for all experiments except SPE comparison and quantification of detection limits. For the SPE comparison experiment, 400 µL of the mix were dried down. For the quantification of metabolite classes, a serial dilution of the mix was prepared to obtain concentrations between 0.5 nmol and 80 nmol of each compound. All dried mixture samples were stored at 4 °C.

### SeaMet metabolite derivatization

To prepare marine samples for gas chromatography-mass spectrometry (GC-MS) analysis, 0.5 to 1 mL of a saltwater sample or experimental mixture dissolved in ASW was dried in a speed vacuum concentrator for 8 hours (Eppendorf Concentrator Plus^(R)^, 45°C, V-AQ). To further remove residual water locked within the salt pellet, 250 µL of toluene (99.8%, < 0.2 % water) was added to each sample and the mixture was ultrasonicated for 10 min at maximum intensity. The toluene was subsequently removed under a gentle flow of N_2_ gas. Metabolite derivatization was performed by adding 80 µL of MeOX dissolved in pyridine (20 mg × mL^-1^) to the dried pellet. The mixture was ultrasonicated (EMag Emmi-12HC®) for 10 min at maximum intensity, briefly vortexed to dissolve the pellet into solution, and subsequently incubated for 90 min at 37 °C using a thermal rotating incubator under constant rotation at 1350 rpm. The pyridine was removed from the sample at room temperature under a gentle flow of N_2_ gas (approximately 1 hour). Following the addition of 100 µL of BSTFA, the mixture was ultrasonicated for 10 min at maximum intensity, vortexed, and incubated for 30 min at 37 °C using a thermal rotating incubator under constant rotation at 1350 rpm. The derivatized mixture was ultrasonicated for 10 min at maximum intensity. Remaining salt in each sample was pelleted through centrifugation at 21.1 *g* for 2 min at 4 °C. 100 µL was transferred to a GC-MS vial for analysis. The full proposed method is publicly available at dx.doi.org/10.17504/protocols.io.nyxdfxn.

### GC-MS data acquisition

All derivatized samples were analyzed on an Agilent 7890B GC coupled to an Agilent 5977A single quadrupole mass selective detector. Using an Agilent 7693 autosampler, 1 µL was injected in splitless mode through a GC inlet liner (ultra inert, splitless, single taper, glass wool, Agilent) onto a DB-5MS column (30 m × 0.25 mm, film thickness 0.25 µm; including 10 m DuraGuard column, Agilent). The inlet liner was changed every 50 samples to avoid damage to the GC column and associated shifts in retention times. The injector temperature was set at 290 °C. Chromatography was achieved with an initial column oven temperature set at 60 °C followed by a ramp of 20 °C min^-1^ until 325 °C, then held for 2 mins. Helium carrier gas was used at a constant flow rate of 1 mL min^-1^. Mass spectra were acquired in electron ionization mode at 70 eV across the mass range of 50–600 m/z and a scan rate of 2 scans s^-1^. The retention time for the method locked using standard mixture of fatty acid methyl esters (Sigma Aldrich).

### Data processing and analysis

Raw Agilent data files were converted to mzXML files using Msconvert (46) and imported into XCMS (v. 2.99.6)(47) within the R software environment (v. 3.4.2) for data processing and analysis. Total ion chromatograms (TIC) were obtained using the xcmsRaw function. TICs comparing sample preparation steps were expressed as a percentage of the MQ control. For environmental and cell culture GC-MS profiles, peaks were picked using the matchedFilter algorithm in XCMS with a full width at half maximum set to 8.4, signal to noise threshold at 1, *m*/*z* width of 0.25 (step parameter), and *m*/*z* difference between overlapping peaks at 1 (**Supplemental Text 1**). Resulting peaks were grouped, retention times corrected and regrouped using the density (bandwidth parameter set to 2) and obiwarp methods. Following peak filling, the CAMERA (v.1.32.0)(48) package was used to place *m*/*z* peaks into pseudo-spectra by grouping similar peaks with the groupFWHM function. Masses below 150 *m*/*z* were removed from the resulting peak table and all profiles were normalized to the ribitol internal standard. Peaks occurring in run blanks and those with higher relative standard deviation scores (% RSD > 25) in quality control samples (cell culture experiment only) were removed from the dataset. To determine differences in metabolite abundances between sediment habitats, metabolite peak data were analyzed using a Bray-Curtis informed non-metric multidimensional scaling analysis followed by an analysis of variance using distance matrices (ADONIS) to test if there are significant differences in metabolite composition between sites. To identify individual peaks that differed significantly between sediment habitats and between cell culture sampling time points, resulting peaks tables were also log transformed and compared using a one-way analysis of variance. All p-values were adjusted using the Benjamini-Hochberg (B.H.) method to control for false positives (49). Significant variables exhibiting large fold-change differences between starting and ending conditions were further investigated. CAMERA grouped peaks from the environmental survey, and those important to shifts in the cell culture experiment were identified using AMDIS (50). Peaks with NIST hits below 800 were compared to the online data repositories, BinVestigate (44) and Golm (43) using the calcualted Kovats retention indices (51) based on a reference n-alkane standard (C7-C40 Saturated Alkanes Standards, Sigme-Aldrich). If no hit was provided, these were considered unknowns.

### The effect of salt and water on metabolite detection

To test the effect of salt on metabolite derivatization, metabolite mix aliquots were resuspended in 1 mL of ASW ranging in salinity from 0 to 34‰ and dried as described above. Methoxamine-trimethylsilylation (TMS) two step derivatization was performed by resuspending each sample in 80 µL of MeOX in pyridine (20 mg mL^-1^) and incubating for 90 min at 37 °C using a thermal rotating incubator under constant rotation at 1350 rpm. MSTFA was subsequently added to the mixture, and the mixture incubated under the same conditions for 90 min (29). Derivatized samples were centrifuged to pellet salt and the supernatant was transferred to a GC-MS vial for analysis. To test the independent effect of water on metabolite derivatization reactions, MQ was added to dried mixture aliquots in steps of 1 µL from 0 to 10 µL. Replicate water gradient samples were subsequently derivatized as before using MeOX and MSTFA or by replacing the MSTFA reagent with BSTFA.

### Marine metabolomics method development

To show how each method development step increased signal intensity and reduced variation in metabolite detection, replicate mixture aliquots (*n* = 5) were resuspended in 0.5 mL of ASW. Mixture aliquots (*n* = 5) were also resuspended in MQ as a no-salt control to highlight the effects of saltwater on metabolite derivatization. 40 µL ribitol (0.2 mM) and 100 µL cholestane (1 mM) were added to each aliquot as internal standards. MQ and ASW samples were first derivatized following the (i) two-step methoxamine-trimethylsilylation (TMS) previously described. Successive steps in the proposed protocol were then applied to ASW samples to demonstrate the combined effects on metabolite detection: (ii) exchange of MSTFA for BSTFA, (iii) removal of residual water from the salt pellet by increasing the speed vacuum drying time and by introducing a toluene drying step to help extract water from the salt pellet, (iv) ultrasonication of the samples after the steps involving addition of toluene, MeOX, BSTFA and following the last derivatization step, and (v) drying the MeOX in pyridine reagent between derivatization reactions. Resulting GC-MS profiles were used to show increases in total signals detected with successive changes in the proposed protocol. Additionally, a cloud plot (using processed peak integration data) was generated to compare compounds dissolved in seawater and to show which metabolite ions exhibited significant (B.H. adjusted *p* < 0.05) and large fold changes (log2(FC) > 2) between the standard and the SeaMet method.

### Solid phase extraction

Replicate metabolite mix aliquots (*n =* 6) were resuspended in 2 mL of artificial seawater. 0.5 mL was reserved from each sample to compare GC-MS profiles before and after SPE sample concentration. Inorganic salts were eluted and metabolites extracted from the remaining 1.5 mL mixture following a SPE based technique using Bond Elut styrene-divinylbenzene (PPL) columns (17). The internal standards ribitol and cholestane were added to both, the reserved sample (before SPE) and the resulting SPE-concentrated sample (after SPE).

All samples were prepared for GC-MS analysis following the proposed marine metabolomics method. Resulting profiles were compared using a cloud plot to show which metabolite ions exhibited significant (B.H. adjusted *p* < 0.05) and large fold changes (log2(FC) > 2) between the pre- and post-SPE treatments.

### Environmental sampling

Replicate porewater profiles were collected from coralline (*n* = 4) and mangrove (*n* = 6) sediments from Carrie Bow Cay (N 16° 04’ 59”, W 88° 04’ 55”) and Twin Cayes, Belize (N 16° 50’ 3”, W 88° 6’ 23”) using a 1 m steel lance with a 2 µm inner diameter covered by 0.063 mm steel mesh. Samples (2 mL water) were collected every 5 cm from the sediment surface to 15 cm depth. Samples were immediately frozen at −20 °C until further analysis. Directly before preparation for GC-MS, the internal standards ribitol and cholestane were added to 0.5 mL of each environmental sample. The mixture was subsequently prepared for GC-MS analysis using the SeaMet method described above.

### Cell culture sampling

Replicate cultures (*n* = 3) *of Marinobacter adhaerens* HP15 DsRed were cultivated in Marine Broth media at 18 °C and 240 rpm as previously described (34). Media samples from the cell cultures and a no-bacteria control media were collected at 0, 4, 8, 12, and 22 h post culture inoculation. Cell counts were monitored at each time point by measuring the optical density at 600 nm (OD_600_). Sampling was carried out by collecting 2 mL of each culture and pelleting the cells through centrifugation for 10 min, at 21.1 *g* and 4 °C. The supernatant was immediately stored at −20 °C until preparation for GC-MS analysis. Prior to sample derivatization using SeaMet, ribitol (0.2 mM; 40 µL) and cholestane (100 mM; 100 µL) were added to 0.5 mL of each experimental sample and subsequently dried down in a speed vacuum concentrator (8 hr, 45 °C, VA-Q). To control for technical variation, quality control (QC) samples (*n* = 3) were prepared by combining 0.25 µL of each culture supernatant and an extraction blank generated by drying down 0.5 mL of MQ.

## Supporting information

Supporting information includes supporting tables, figures, references and XCMS peak picking script.

## Acknowledgements

We thank T. Gulstad, K. Caspersen, F. Fojt, and M. Meyer (MPI-Bremen) for support with data acquisition and sample preparation, N. Böttcher (Jacobs University Bremen) for providing culture samples, D. Michellod for sediment pore water sample collection, and B. Geier and J. Beckmann (MPI-Bremen) for valuable discussions. We acknowledge the Max-Planck Society and the Gordon and Betty Moore Foundation (Marine Microbial Initiative Investigator Award to ND, Grant #GBMF3811) for financial support. This work is contribution *XXX* from the Carrie Bow Cay Laboratory, Caribbean Coral Reef Ecosystem Program, National Museum of Natural History, Washington DC.

## Supplementary Information for

### Supporting Figures and Tables

**Figure S1.**
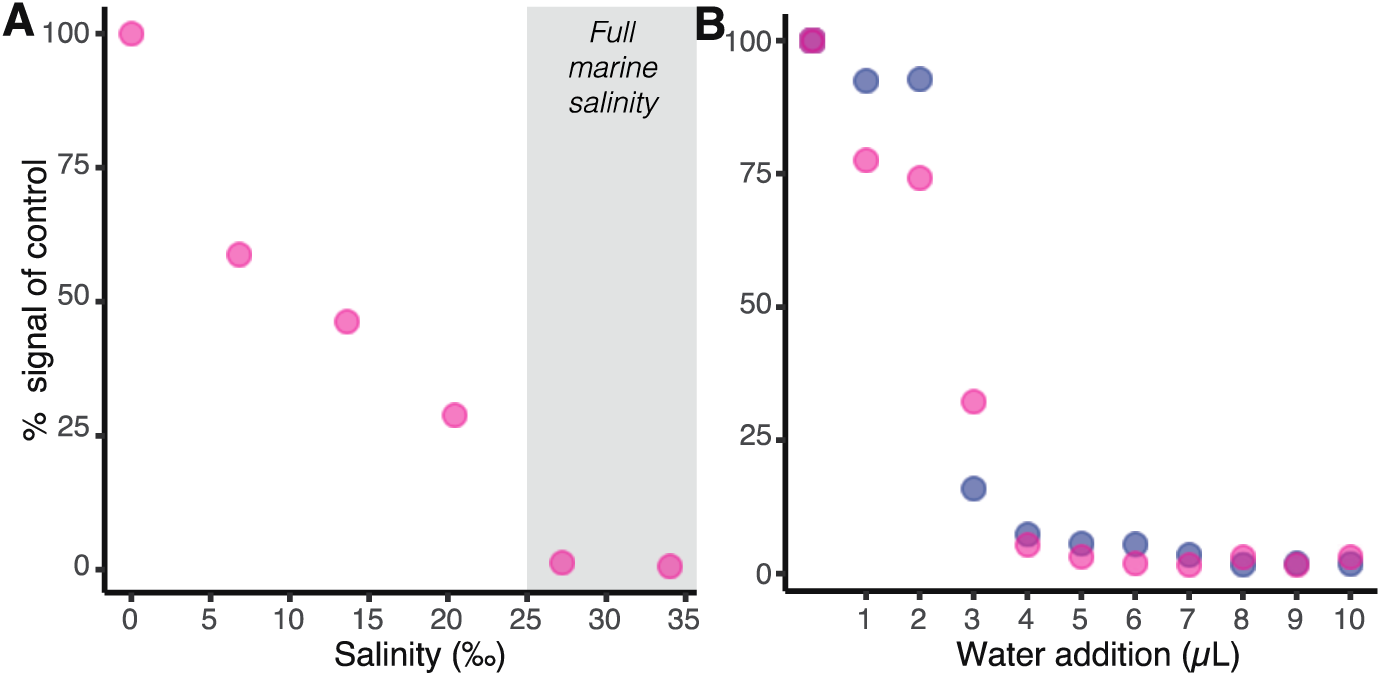
Salt and water inhibit metabolite derivatization reactions. Total detected ion signals are negatively related to increasing concentrations of **A**, salt and **B**, water for both MSTFA (red circles) and BSTFA (blue circles) derivatization reagents. Signals intensities from the metabolite mixture (**Table S6**) are relative to control samples (**A**, no salt; **B**, no water). To avoid damage to the GC-MS instrumentation, only 1 replicate / condition was analyzed.

**Figure S2.**
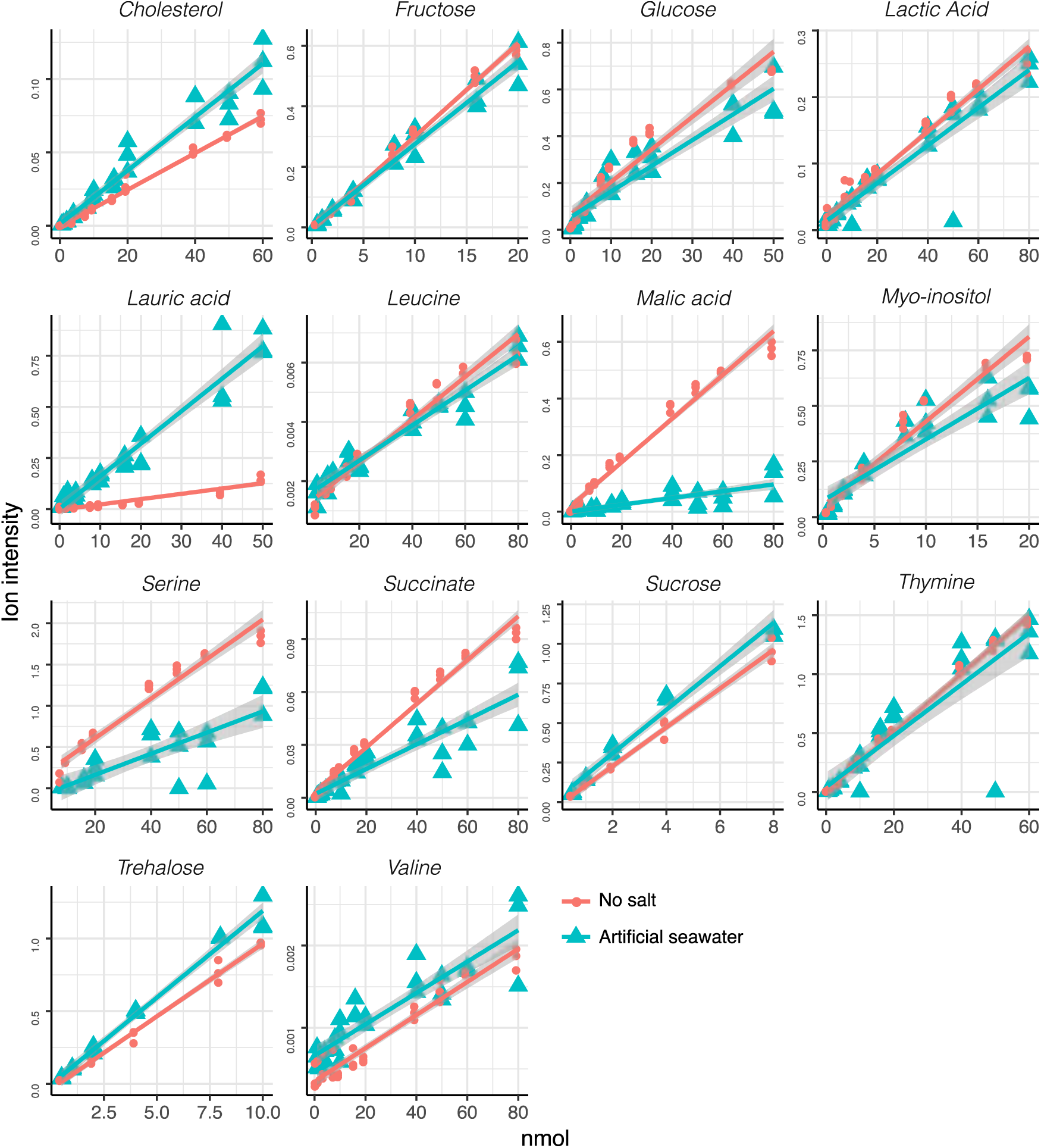
Calibration curves for individual metabolites in salt free and artificial seawater (ASW). Calculated calibration curves were compared for compounds that were detected in both salt-free and ASW conditions (n=3 for each concentration). Gray shading represents 90% confidence intervals and points are fitted using a linear regression. Model results are reported in Table S2.

**Figure S3.**
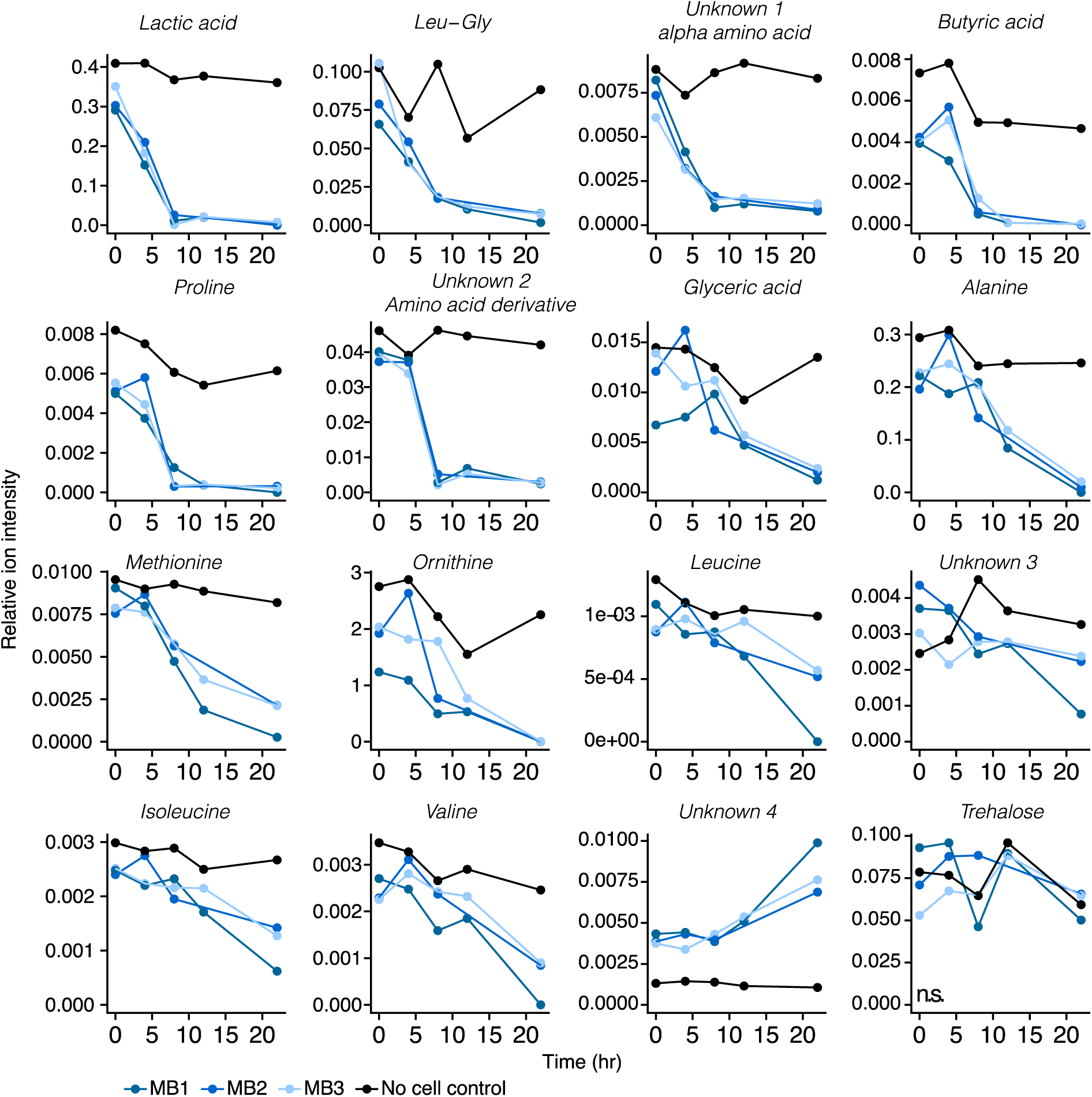
Extracellular metabolite levels shift with cell culture density. Metabolite relative abundances for each cell culture and no-cell control are plotted through time. Only metabolites that significantly (adjusted *p-*value < 0.05) varied with time in replicate culture experiments are plotted for clarity.

**Table S1.**
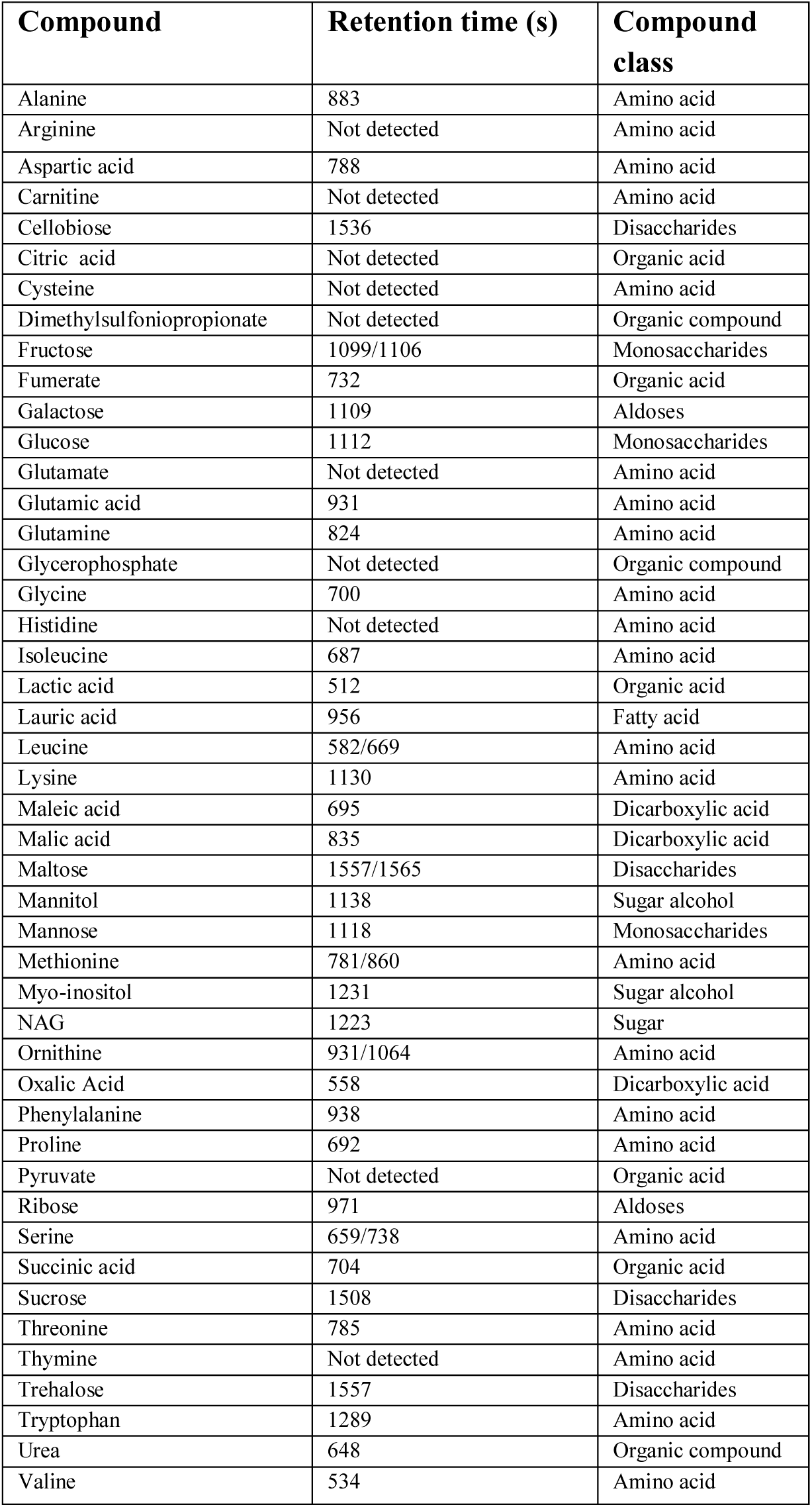
Mixture of 45 metabolites used for the development and testing of the sample preparation method, with their retention times as detected in samples dissolved in artificial seawater using GC-MS. Metabolites with multiple retention times represent different TMS derivatives.

**Table S2.** Quantification ions, calibration coefficients, and retention of metabolites from each compound class in no salt and salt conditions. The minimum and maximum calibration points of select compounds from representatives of each metabolite class in artificial seawater are reported. The lowest concentration at which signals were observed is reported for select compounds.

| Compound | Category | quantification<br>ion (m/z) | No salt |  |  |  | Salt |  |  |  | Lowest<br>conc.** |
| --- | --- | --- | --- | --- | --- | --- | --- | --- | --- | --- | --- |
|  |  |  | Retention<br>time (s) | r<sup>2</sup> | %CV<br>4<br>nmol | Retention<br>time (s) | r<sup>2</sup> | %<br>CV<br>4<br>nmol | Min<br>calibration<br>point<br>(nmol)* | Max<br>calibration<br>point<br>(nmol)* |  |
| Glycine 3TMS | Amino acid | 204.1 | 689 | 0.955 | 17.12 | 689 | < 0.7 | 4.68 | 0.5 | 80 |  |
| Histidine 2TMS | Amino acid | 218.1 | 1126 | 0.973 | 20.73 | Not detected |  |  |  |  |  |
| Leucine 2 TMS | Amino acid | 177.06 | 659 | 0.961 | 17.24 | 660 | 0.939 | 25.93 | 4 | 80 |  |
| Lysine 3TMS | Amino acid | 156.1 | 989 | 0.973 | 31.31 | Not detected |  |  |  |  |  |
| Lysine 4TMS | Amino acid | 174.1 | Not detected |  |  | 1127 | < 0.7 | 15.28 | 0.5 | 80 |  |
| Ornithine 3TMS | Amino acid | 174.1 | 928 | 0.992 | 18.19 | Not detected |  |  |  |  |  |
| Ornithine 4TMS | Amino acid | 174.1 | Not detected |  |  | 1061 | < 0.7 | 17.38 | 0.5 | 80 |  |
| Serine 3TMS | Amino acid | 204.16 | 732 | 0.957 | NA | 733 | < 0.7 |  | 8 | 80 |  |
| Thymine 2TMS | Amino acid | 255.1 | 765 | 0.996 | 22.65 | 766 | 0.775 | 7.11 | 0.5 | 60 | 0.25 |
| Valine 2TMS | Amino acid | 188.09 | 612 | 0.94 | 32.15 | 614 | 0.828 | 10.94 | 1 | 80 | 0.25 |
| Citrate/citric acid 3TMS | Organic acid | 244.14 | 1061 | 0.954 | 16.68 | Not detected |  |  |  |  |  |
| Glutamic acid 3TMS | Organic acid | 339.13 | 933 | < 0.7 | 46.64 | Not detected |  |  |  |  |  |
| Glutamic acid 3TMS | Organic acid | 246.1 | Not detected |  |  | 929 | < 0.7 | 18.1 | 1 | 80 |  |
| Glutamic acid dervative - Hydroxproline | Organic acid | 216.09 | 860 | 0.995 | 18.33 | Not detected |  |  |  |  |  |
| Glutamic acid dervative - Hydroxproline | Organic acid | 221.09 | 860 | < 0.7 | 14.13 | 859 | < 0.7 | 4.68 | 0.5 | 80 |  |
| Lactic Acid 2TMS | Organic acid | 191.1 | 478 | 0.971 | 20.21 | 480 | 0.877 | 7.28 | 0.5 | 80 | 0.0312 |
| Lauric acid TMS | Organic acid | 257.2 | 955 | 0.903 | 34.18 | 955 | 0.938 | 37.16 | 0.5 | 50 | 0.0312 |
| Malic acid 3TMS | Organic acid | 233.1 | 832 | 0.98 | 14.04 | 832 | < 0.7 | 11.45 | 0.5 | 80 |  |
| Succinate 2TMS | Organic acid | 172.1 | 696 | 0.982 | 15.94 | 696 | 0.835 | 5.71 | 0.5 | 80 | 0.25 |
| Cholesterol TMS | Sterol | 247.25 | 1716 | 0.99 | 34.42 | 1717 | 0.954 | 26.18 | 0.5 | 60 | 0.5 |
| Fructose 5TMS | Sugar | 307.2 | 1093 | 0.994 | 12.6 | 1093 | 0.971 | 16.89 | 0.5 | 20 | 0.0312 |
| Glucose 5TMS | Sugar | 205.1 | 1121 | 0.982 | 24.61 | 1120 | 0.804 | 15.18 | 0.5 | 80 | 0.125 |
| Sucrose 8TMS | Sugar | 361.27 | 1495 | 0.989 | 13.5 | 1495 | 0.975 | 2.44 | 0.5 | 8 | 0.0312 |
| **Trehalose**<br>**8TMS** | Sugar | 361.27 | 1543 | 0.991 | 12.91 | 1543 | 0.985 | 2.07 | 0.5 | 10 | 0.0312 |
| **Myo-inositol**<br>**6TMS** | Sugar<br>alcohol | 305 | 1223 | 0.947 | 14.19 | 1224 | 0.853 | 14.34 | 0.5 | 20 | 0.0312 |

CV = % coefficient of variation \*nmol in 0.5 mL of artificial seawater \*\* For selected compounds in artificial seawater

**Table S3.**
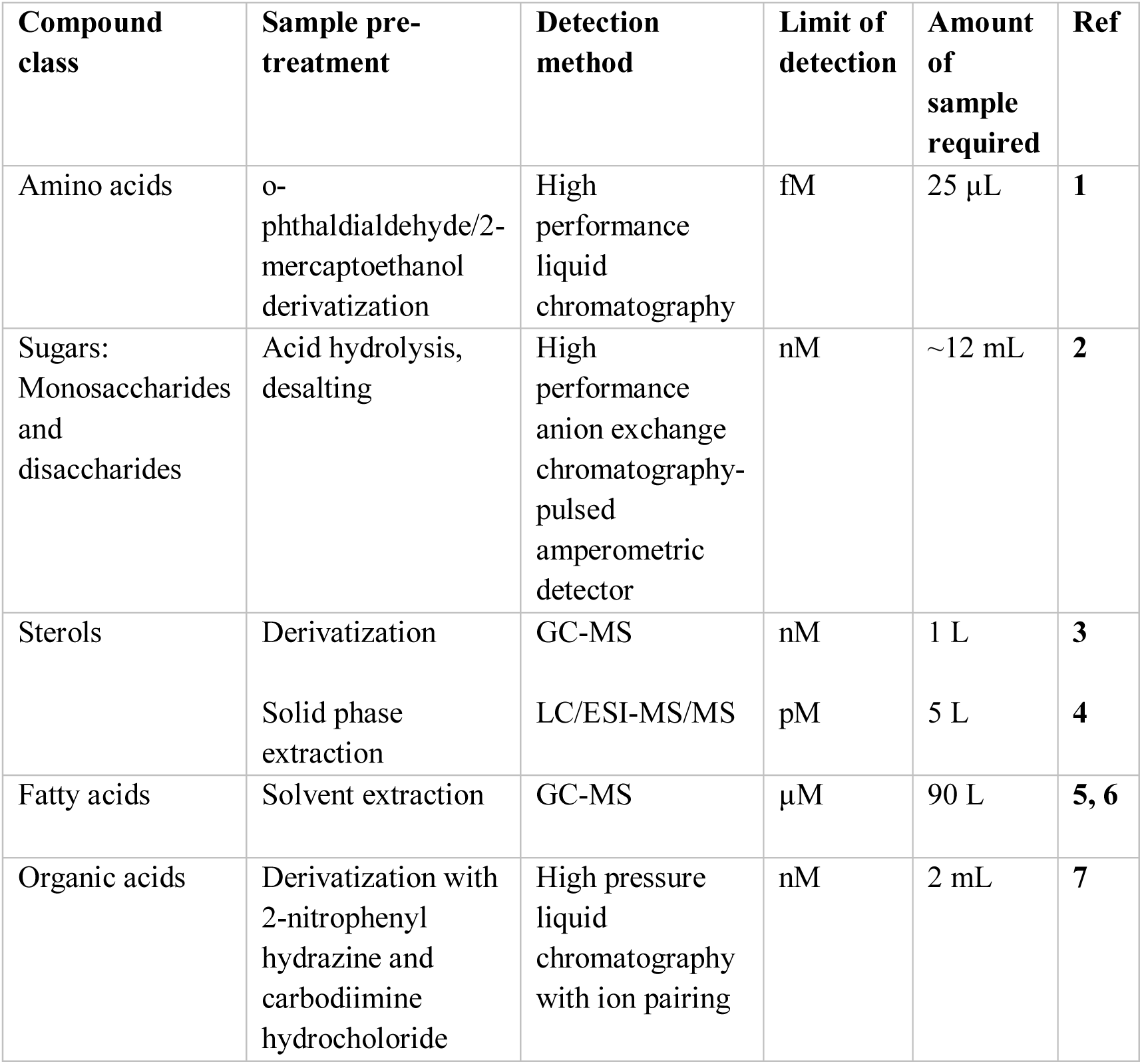
Targeted techniques for the quantification of specific metabolite classes in seawater.

**Table S4.**
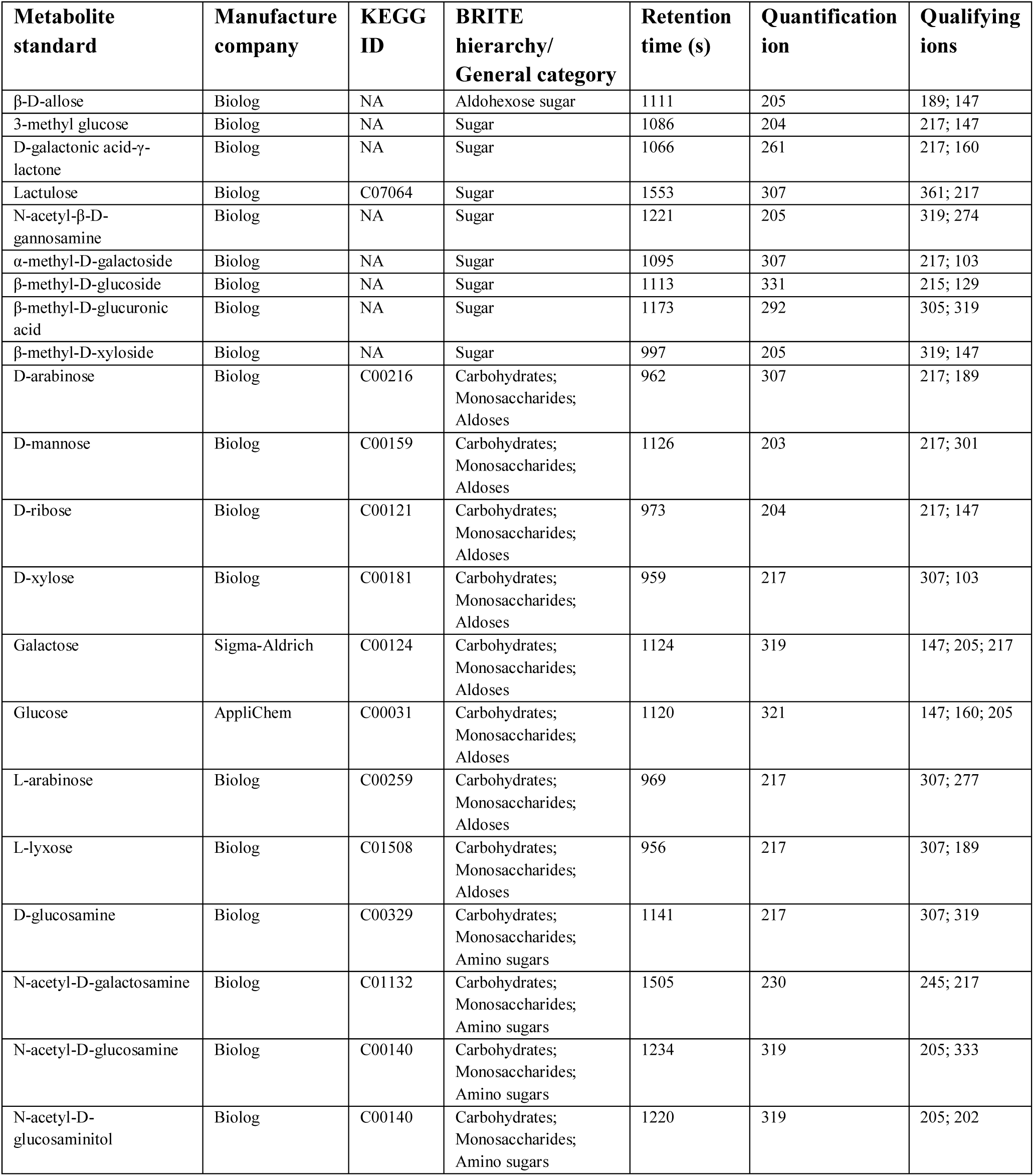
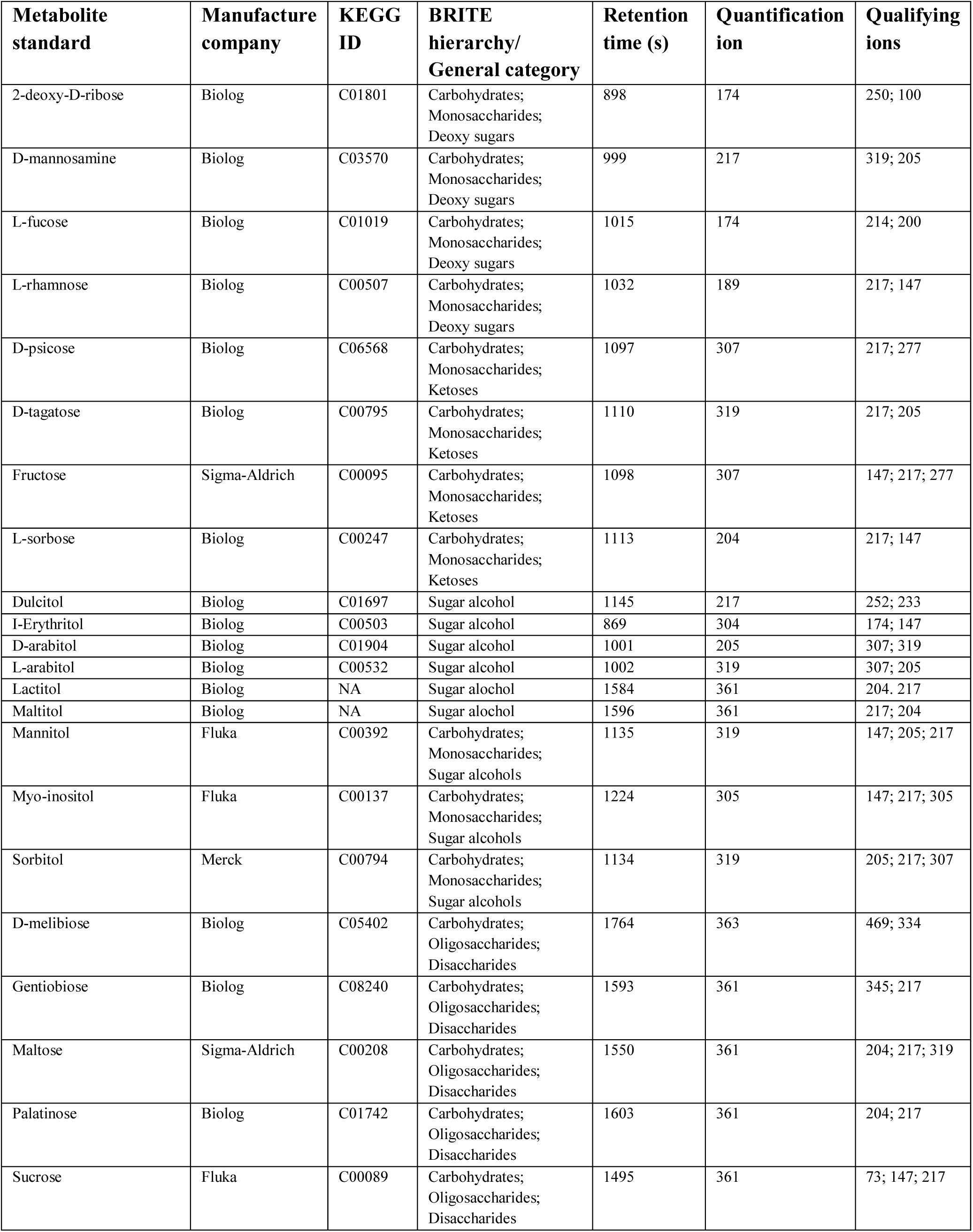
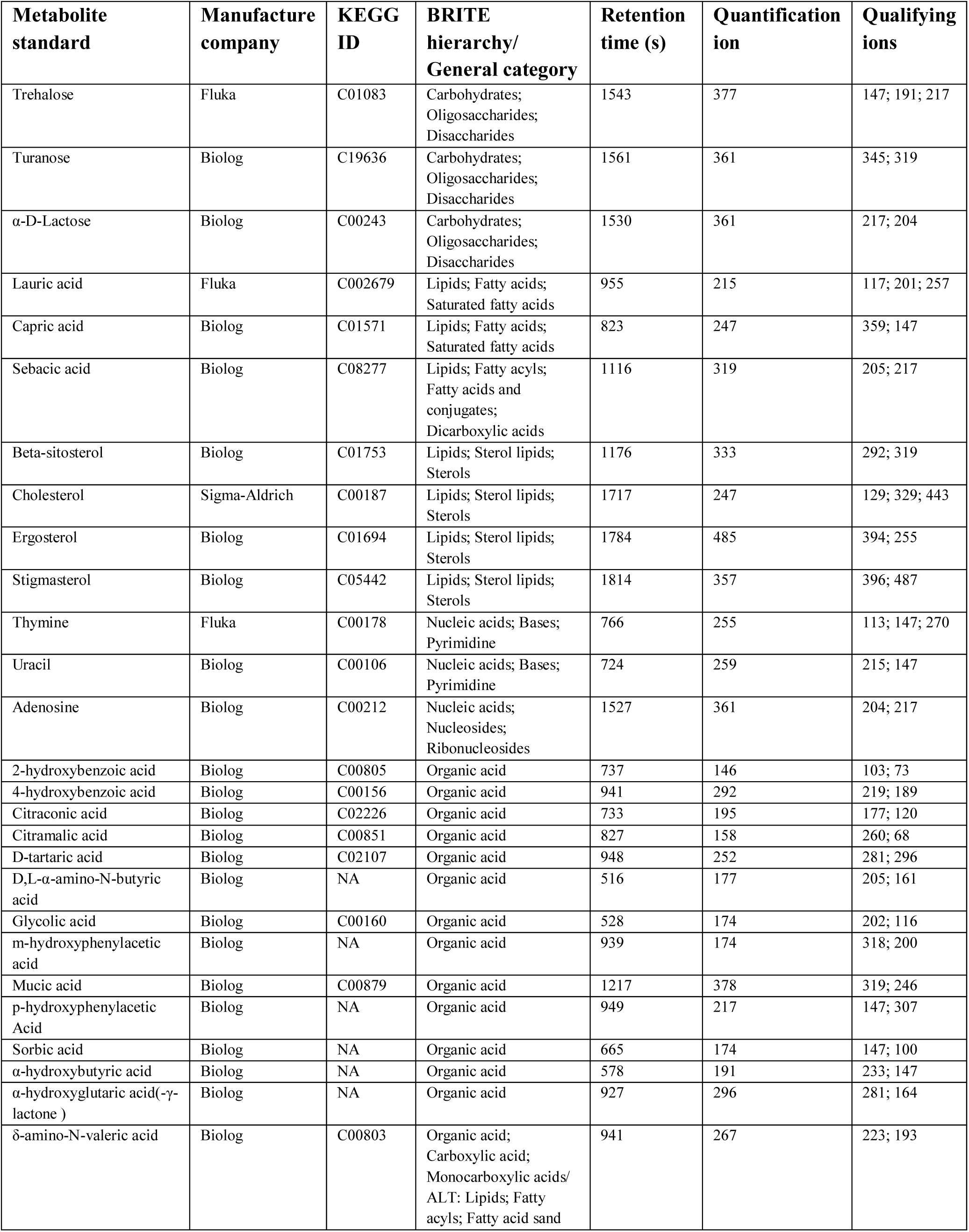
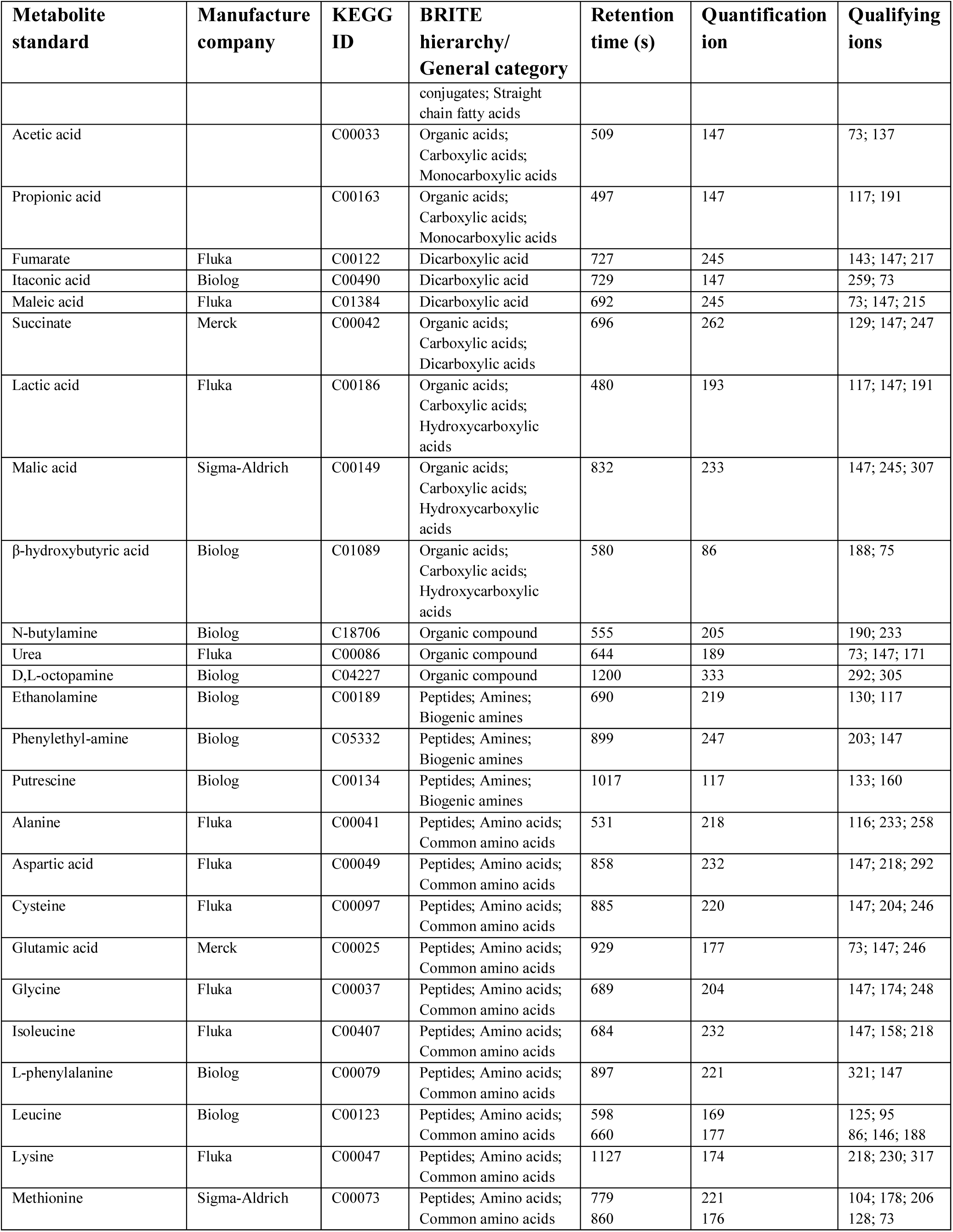
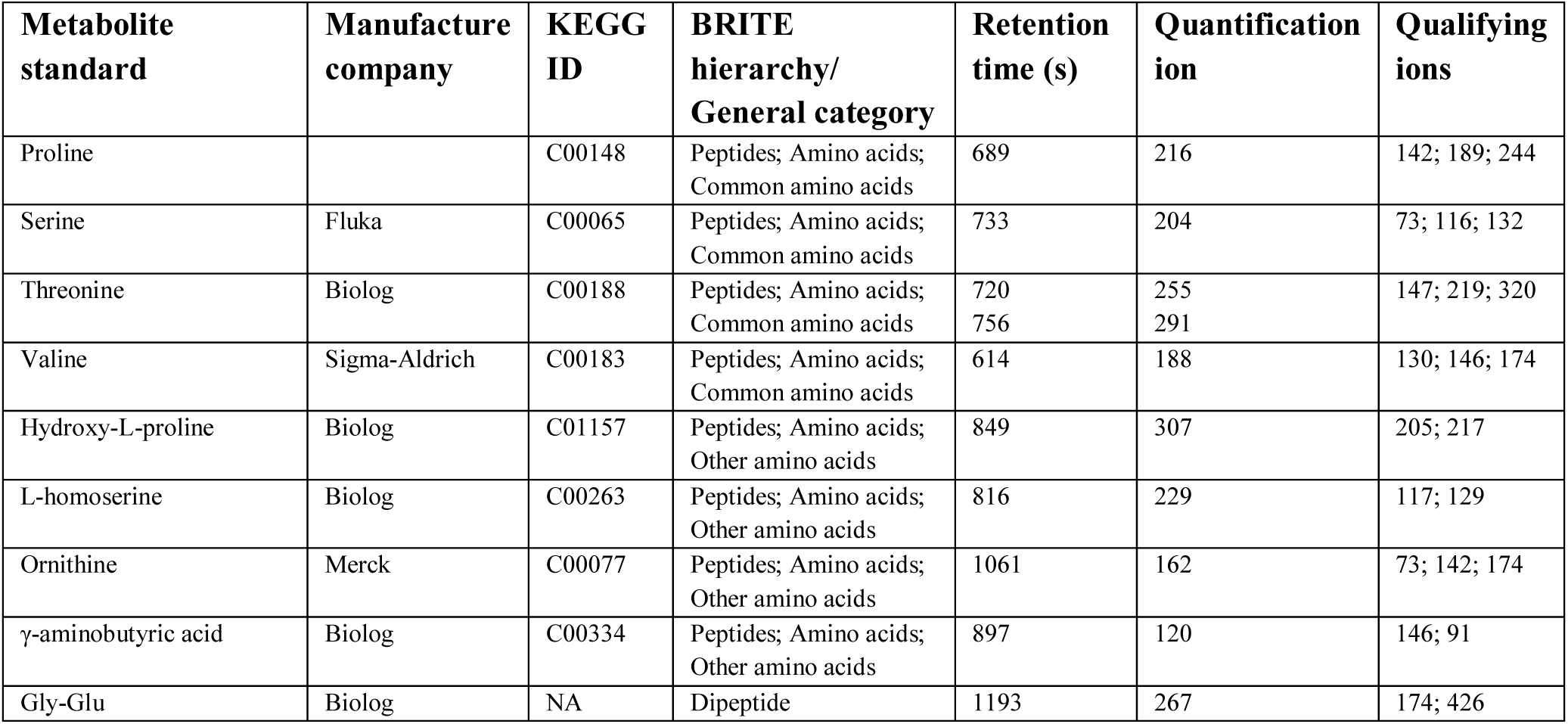
Retention times, quantification and qualification ions of the 107 metabolite standards detected in artificial seawater using SeaMet. Kyoto Encyclopedia of Genes and Genomes (KEGG ID) ids and associated BRITE functional hierarchy or metabolite class of each compound are indicated.

| Metabolite standard | Manufacture company | KEGG ID | BRITE hierarchy/<br>General category | Retention time (s) | Quantification ion | Qualifying ions |
| --- | --- | --- | --- | --- | --- | --- |
| β-D-allose | Biolog | NA | Aldohexose sugar | 1111 | 205 | 189; 147 |
| 3-methyl glucose | Biolog | NA | Sugar | 1086 | 204 | 217; 147 |
| D-galactonic acid-γ-lactone | Biolog | NA | Sugar | 1066 | 261 | 217; 160 |
| Lactulose | Biolog | C07064 | Sugar | 1553 | 307 | 361; 217 |
| N-acetyl-β-D-galactosamine | Biolog | NA | Sugar | 1221 | 205 | 319; 274 |
| α-methyl-D-galactoside | Biolog | NA | Sugar | 1095 | 307 | 217; 103 |
| β-methyl-D-glucoside | Biolog | NA | Sugar | 1113 | 331 | 215; 129 |
| β-methyl-D-glucuronic acid | Biolog | NA | Sugar | 1173 | 292 | 305; 319 |
| β-methyl-D-xyloside | Biolog | NA | Sugar | 997 | 205 | 319; 147 |
| D-arabinose | Biolog | C00216 | Carbohydrates;<br>Monosaccharides;<br>Aldoses | 962 | 307 | 217; 189 |
| D-mannose | Biolog | C00159 | Carbohydrates;<br>Monosaccharides;<br>Aldoses | 1126 | 203 | 217; 301 |
| D-ribose | Biolog | C00121 | Carbohydrates;<br>Monosaccharides;<br>Aldoses | 973 | 204 | 217; 147 |
| D-xylose | Biolog | C00181 | Carbohydrates;<br>Monosaccharides;<br>Aldoses | 959 | 217 | 307; 103 |
| Galactose | Sigma-Aldrich | C00124 | Carbohydrates;<br>Monosaccharides;<br>Aldoses | 1124 | 319 | 147; 205; 217 |
| Glucose | AppliChem | C00031 | Carbohydrates;<br>Monosaccharides;<br>Aldoses | 1120 | 321 | 147; 160; 205 |
| L-arabinose | Biolog | C00259 | Carbohydrates;<br>Monosaccharides;<br>Aldoses | 969 | 217 | 307; 277 |
| L-lyxose | Biolog | C01508 | Carbohydrates;<br>Monosaccharides;<br>Aldoses | 956 | 217 | 307; 189 |
| D-glucosamine | Biolog | C00329 | Carbohydrates;<br>Monosaccharides;<br>Amino sugars | 1141 | 217 | 307; 319 |
| N-acetyl-D-galactosamine | Biolog | C01132 | Carbohydrates;<br>Monosaccharides;<br>Amino sugars | 1505 | 230 | 245; 217 |
| N-acetyl-D-glucosamine | Biolog | C00140 | Carbohydrates;<br>Monosaccharides;<br>Amino sugars | 1234 | 319 | 205; 333 |
| N-acetyl-D-glucosaminitol | Biolog | C00140 | Carbohydrates;<br>Monosaccharides;<br>Amino sugars | 1220 | 319 | 205; 202 |

| **Metabolite standard** | **Manufacture company** | **KEGG ID** | **BRITE hierarchy/**<br>**General category** | **Retention time (s)** | **Quantification ion** | **Qualifying ions** |
| --- | --- | --- | --- | --- | --- | --- |
| 2-deoxy-D-ribose | Biolog | C01801 | Carbohydrates;<br>Monosaccharides;<br>Deoxy sugars | 898 | 174 | 250; 100 |
| D-mannosamine | Biolog | C03570 | Carbohydrates;<br>Monosaccharides;<br>Deoxy sugars | 999 | 217 | 319; 205 |
| L-fucose | Biolog | C01019 | Carbohydrates;<br>Monosaccharides;<br>Deoxy sugars | 1015 | 174 | 214; 200 |
| L-rhamnose | Biolog | C00507 | Carbohydrates;<br>Monosaccharides;<br>Deoxy sugars | 1032 | 189 | 217; 147 |
| D-psicose | Biolog | C06568 | Carbohydrates;<br>Monosaccharides;<br>Ketoses | 1097 | 307 | 217; 277 |
| D-tagatose | Biolog | C00795 | Carbohydrates;<br>Monosaccharides;<br>Ketoses | 1110 | 319 | 217; 205 |
| Fructose | Sigma-Aldrich | C00095 | Carbohydrates;<br>Monosaccharides;<br>Ketoses | 1098 | 307 | 147; 217; 277 |
| L-sorbose | Biolog | C00247 | Carbohydrates;<br>Monosaccharides;<br>Ketoses | 1113 | 204 | 217; 147 |
| Dulcitol | Biolog | C01697 | Sugar alcohol | 1145 | 217 | 252; 233 |
| L-Erythritol | Biolog | C00503 | Sugar alcohol | 869 | 304 | 174; 147 |
| D-arabitol | Biolog | C01904 | Sugar alcohol | 1001 | 205 | 307; 319 |
| L-arabitol | Biolog | C00532 | Sugar alcohol | 1002 | 319 | 307; 205 |
| Lactitol | Biolog | NA | Sugar alcohol | 1584 | 361 | 204; 217 |
| Maltitol | Biolog | NA | Sugar alcohol | 1596 | 361 | 217; 204 |
| Mannitol | Fluka | C00392 | Carbohydrates;<br>Monosaccharides;<br>Sugar alcohols | 1135 | 319 | 147; 205; 217 |
| Myo-inositol | Fluka | C00137 | Carbohydrates;<br>Monosaccharides;<br>Sugar alcohols | 1224 | 305 | 147; 217; 305 |
| Sorbitol | Merck | C00794 | Carbohydrates;<br>Monosaccharides;<br>Sugar alcohols | 1134 | 319 | 205; 217; 307 |
| D-melibiose | Biolog | C05402 | Carbohydrates;<br>Oligosaccharides;<br>Disaccharides | 1764 | 363 | 469; 334 |
| Gentiobiose | Biolog | C08240 | Carbohydrates;<br>Oligosaccharides;<br>Disaccharides | 1593 | 361 | 345; 217 |
| Maltose | Sigma-Aldrich | C00208 | Carbohydrates;<br>Oligosaccharides;<br>Disaccharides | 1550 | 361 | 204; 217; 319 |
| Palatinose | Biolog | C01742 | Carbohydrates;<br>Oligosaccharides;<br>Disaccharides | 1603 | 361 | 204; 217 |
| Sucrose | Fluka | C00089 | Carbohydrates;<br>Oligosaccharides;<br>Disaccharides | 1495 | 361 | 73; 147; 217 |
| Trehalose | Fluka | C01083 | Carbohydrates;<br>Oligosaccharides;<br>Disaccharides | 1543 | 377 | 147; 191; 217 |
| Turanose | Biolog | C19636 | Carbohydrates;<br>Oligosaccharides;<br>Disaccharides | 1561 | 361 | 345; 319 |
| α-D-Lactose | Biolog | C00243 | Carbohydrates;<br>Oligosaccharides;<br>Disaccharides | 1530 | 361 | 217; 204 |
| Lauric acid | Fluka | C002679 | Lipids; Fatty acids;<br>Saturated fatty acids | 955 | 215 | 117; 201; 257 |
| Capric acid | Biolog | C01571 | Lipids; Fatty acids;<br>Saturated fatty acids | 823 | 247 | 359; 147 |
| Sebacic acid | Biolog | C08277 | Lipids; Fatty acyls;<br>Fatty acids and<br>conjugates;<br>Dicarboxylic acids | 1116 | 319 | 205; 217 |
| Beta-sitosterol | Biolog | C01753 | Lipids; Sterol lipids;<br>Sterols | 1176 | 333 | 292; 319 |
| Cholesterol | Sigma-Aldrich | C00187 | Lipids; Sterol lipids;<br>Sterols | 1717 | 247 | 129; 329; 443 |
| Ergosterol | Biolog | C01694 | Lipids; Sterol lipids;<br>Sterols | 1784 | 485 | 394; 255 |
| Stigmasterol | Biolog | C05442 | Lipids; Sterol lipids;<br>Sterols | 1814 | 357 | 396; 487 |
| Thymine | Fluka | C00178 | Nucleic acids; Bases;<br>Pyrimidine | 766 | 255 | 113; 147; 270 |
| Uracil | Biolog | C00106 | Nucleic acids; Bases;<br>Pyrimidine | 724 | 259 | 215; 147 |
| Adenosine | Biolog | C00212 | Nucleic acids;<br>Nucleosides;<br>Ribonucleosides | 1527 | 361 | 204; 217 |
| 2-hydroxybenzoic acid | Biolog | C00805 | Organic acid | 737 | 146 | 103; 73 |
| 4-hydroxybenzoic acid | Biolog | C00156 | Organic acid | 941 | 292 | 219; 189 |
| Citraconic acid | Biolog | C02226 | Organic acid | 733 | 195 | 177; 120 |
| Citramalic acid | Biolog | C00851 | Organic acid | 827 | 158 | 260; 68 |
| D-tartaric acid | Biolog | C02107 | Organic acid | 948 | 252 | 281; 296 |
| D,L-α-amino-N-butyric acid | Biolog | NA | Organic acid | 516 | 177 | 205; 161 |
| Glycolic acid | Biolog | C00160 | Organic acid | 528 | 174 | 202; 116 |
| m-hydroxyphenylacetic acid | Biolog | NA | Organic acid | 939 | 174 | 318; 200 |
| Mucic acid | Biolog | C00879 | Organic acid | 1217 | 378 | 319; 246 |
| p-hydroxyphenylacetic Acid | Biolog | NA | Organic acid | 949 | 217 | 147; 307 |
| Sorbic acid | Biolog | NA | Organic acid | 665 | 174 | 147; 100 |
| α-hydroxybutyric acid | Biolog | NA | Organic acid | 578 | 191 | 233; 147 |
| α-hydroxyglutaric acid(-γ-lactone ) | Biolog | NA | Organic acid | 927 | 296 | 281; 164 |
| δ-amino-N-valeric acid | Biolog | C00803 | Organic acid;<br>Carboxylic acid;<br>Monocarboxylic acids/<br>ALT: Lipids; Fatty<br>acyls; Fatty acid sand conjugates; Straight chain fatty acids | 941 | 267 | 223; 193 |
| Acetic acid |  | C00033 | Organic acids;<br>Carboxylic acids;<br>Monocarboxylic acids | 509 | 147 | 73; 137 |
| Propionic acid |  | C00163 | Organic acids;<br>Carboxylic acids;<br>Monocarboxylic acids | 497 | 147 | 117; 191 |
| Fumarate | Fluka | C00122 | Dicarboxylic acid | 727 | 245 | 143; 147; 217 |
| Itaconic acid | Biolog | C00490 | Dicarboxylic acid | 729 | 147 | 259; 73 |
| Maleic acid | Fluka | C01384 | Dicarboxylic acid | 692 | 245 | 73; 147; 215 |
| Succinate | Merck | C00042 | Organic acids;<br>Carboxylic acids;<br>Dicarboxylic acids | 696 | 262 | 129; 147; 247 |
| Lactic acid | Fluka | C00186 | Organic acids;<br>Carboxylic acids;<br>Hydroxycarboxylic acids | 480 | 193 | 117; 147; 191 |
| Malic acid | Sigma-Aldrich | C00149 | Organic acids;<br>Carboxylic acids;<br>Hydroxycarboxylic acids | 832 | 233 | 147; 245; 307 |
| β-hydroxybutyric acid | Biolog | C01089 | Organic acids;<br>Carboxylic acids;<br>Hydroxycarboxylic acids | 580 | 86 | 188; 75 |
| N-butylamine | Biolog | C18706 | Organic compound | 555 | 205 | 190; 233 |
| Urea | Fluka | C00086 | Organic compound | 644 | 189 | 73; 147; 171 |
| D,L-octopamine | Biolog | C04227 | Organic compound | 1200 | 333 | 292; 305 |
| Ethanolamine | Biolog | C00189 | Peptides; Amines;<br>Biogenic amines | 690 | 219 | 130; 117 |
| Phenylethyl-amine | Biolog | C05332 | Peptides; Amines;<br>Biogenic amines | 899 | 247 | 203; 147 |
| Putrescine | Biolog | C00134 | Peptides; Amines;<br>Biogenic amines | 1017 | 117 | 133; 160 |
| Alanine | Fluka | C00041 | Peptides; Amino acids;<br>Common amino acids | 531 | 218 | 116; 233; 258 |
| Aspartic acid | Fluka | C00049 | Peptides; Amino acids;<br>Common amino acids | 858 | 232 | 147; 218; 292 |
| Cysteine | Fluka | C00097 | Peptides; Amino acids;<br>Common amino acids | 885 | 220 | 147; 204; 246 |
| Glutamic acid | Merck | C00025 | Peptides; Amino acids;<br>Common amino acids | 929 | 177 | 73; 147; 246 |
| Glycine | Fluka | C00037 | Peptides; Amino acids;<br>Common amino acids | 689 | 204 | 147; 174; 248 |
| Isoleucine | Fluka | C00407 | Peptides; Amino acids;<br>Common amino acids | 684 | 232 | 147; 158; 218 |
| L-phenylalanine | Biolog | C00079 | Peptides; Amino acids;<br>Common amino acids | 897 | 221 | 321; 147 |
| Leucine | Biolog | C00123 | Peptides; Amino acids;<br>Common amino acids | 598<br>660 | 169<br>177 | 125; 95<br>86; 146; 188 |
| Lysine | Fluka | C00047 | Peptides; Amino acids;<br>Common amino acids | 1127 | 174 | 218; 230; 317 |
| Methionine | Sigma-Aldrich | C00073 | Peptides; Amino acids;<br>Common amino acids | 779<br>860 | 221<br>176 | 104; 178; 206<br>128; 73 |
| Proline |  | C00148 | Peptides; Amino acids;<br>Common amino acids | 689 | 216 | 142; 189; 244 |
| Serine | Fluka | C00065 | Peptides; Amino acids;<br>Common amino acids | 733 | 204 | 73; 116; 132 |
| Threonine | Biolog | C00188 | Peptides; Amino acids;<br>Common amino acids | 720<br>756 | 255<br>291 | 147; 219; 320 |
| Valine | Sigma-Aldrich | C00183 | Peptides; Amino acids;<br>Common amino acids | 614 | 188 | 130; 146; 174 |
| Hydroxy-L-proline | Biolog | C01157 | Peptides; Amino acids;<br>Other amino acids | 849 | 307 | 205; 217 |
| L-homoserine | Biolog | C00263 | Peptides; Amino acids;<br>Other amino acids | 816 | 229 | 117; 129 |
| Ornithine | Merck | C00077 | Peptides; Amino acids;<br>Other amino acids | 1061 | 162 | 73; 142; 174 |
| γ-aminobutyric acid | Biolog | C00334 | Peptides; Amino acids;<br>Other amino acids | 897 | 120 | 146; 91 |
| Gly-Glu | Biolog | NA | Dipeptide | 1193 | 267 | 174; 426 |

**Table S5.** Major metabolite peaks found in marine sediment porewaters and their retention times. Compounds that did not match NIST database enteries are labeled as “unknown” followed by their retention time. Mass spectra from these compounds were also compared to both the Golm and BinBase databases using the Kovats retention time index adjustment. Golm predicted functional groups and BinBase splash id’s are reported when unknowns matched database hits.

| Habitat | Annotation | Retention time (s) | Class | Unknown Compounds |  |  |
| --- | --- | --- | --- | --- | --- | --- |
|  |  |  |  | Kovats retention index | Golm predicted functional groups | BinBase related splash ID |
| Coralline | Lactic acid | 499 | Organic acid |  |  |  |
|  | Acetate | 512 | Organic acid |  |  |  |
|  | Alanine | 533 | Amino acid |  |  |  |
|  | Glycerol | 666 | Sugar alcohol |  |  |  |
|  | Glycine | 695 | Amino acid |  |  |  |
|  | Succinate | 699 | Organic acid |  |  |  |
|  | Propanoic acid | 713 | Organic acid |  |  |  |
|  | Unknown 893 | 895 |  | 1563 | No hits | No hits |
|  | Unknown 1031 | 1031 |  | 1759 | Primary alcohol<br>Secondary alcohol<br>Alcohol<br>1,2, diol<br>Hydroxy | splash10-0fvj-0920000000-2b50a374508a5be92557 |
|  | Azelaic acid | 1048 | Fatty Acid |  |  |  |
|  | Unknown compound 1073 | 1073 |  | 1823 | Hydroxy<br>Alcohol<br>Carboxylic acid | No hits |
|  | Pinitol | 1089 | Sugar alcohol |  |  |  |
|  | Fructose | 1101 | Sugar |  |  |  |
|  | Galactose | 1108 | Sugar |  |  |  |
|  | Galactose | 1112 | Sugar |  |  |  |
|  | Mannitol | 1138 | Sugar alcohol |  |  |  |
|  | Sucrose | 1503 | Sugar |  |  |  |
|  | Trehalose | 1553 | Sugar |  |  |  |
| Mangrove | Lactic acid | 499 | Organic acid |  |  |  |
|  | Acetate | 512 | Organic acid |  |  |  |
|  | Glycerol | 666 | Sugar alcohol |  |  |  |
|  | Succinate | 699 | Organic acid |  |  |  |
|  | Pentanoic acid | 914 | Organic acid |  |  |  |
|  | Lauric acid | 937 | Straight chain fatty acid |  |  |  |
|  | Lauric acid | 956 | Straight chain fatty acid |  |  |  |
|  | Unknown 1031 | 1031 |  | 1759 | Primary alcohol<br>Secondary alcohol<br>Alcohol<br>1,2, diol<br>Hydroxy | splash10-0fvj-0920000000-2b50a374508a5be92557 |
|  | Azelaic acid | 1048 | Fatty Acid |  |  |  |
|  | Aromatic dione | 1126 |  |  |  |  |
|  | Mannitol | 1138 | Sugar alcohol |  |  |  |
|  | Palmitic acid | 1177 | Straight chain fatty acid |  |  |  |
|  | Isooctyl laurate | 1235 | Straight chain fatty acid |  |  |  |
|  | Lauric acid | 1289 | Straight chain fatty acid |  |  |  |
|  | Myristic acid | 1371 | Straight chain fatty acid |  |  |  |
|  | Myristic acid | 1388 | Straight chain fatty acid |  |  |  |
|  |  |  |  | Kovats retention index | Golm predicted functional groups | BinBase related splash ID |
|  | Pentadecanoic acid | 1434 | Straight chain fatty acid |  |  |  |
|  | Hexadecanoic acid | 1464 | Straight chain fatty acid |  |  |  |
|  | Hexadecanoic acid | 1481 | Straight chain fatty acid |  |  |  |
|  | Sucrose | 1503 | Sugar |  |  |  |
|  | Heptadecanoic acid | 1509 | Straight chain fatty acid |  |  |  |
|  | Heptadecanoic acid | 1524 | Straight chain fatty acid |  |  |  |
|  | 2-Monostearin | 1552 | Straight chain fatty acid |  |  |  |
|  | Octadecanoic acid | 1569 | Straight chain fatty acid |  |  |  |
|  | Unknown peak 1595 | 1595 |  | 2832 | Alcohol | No hits |
|  | Nonadecanoic acid | 1607 | Straight chain fatty acid |  |  |  |
|  | Unknown 1634 | 1635 |  | 2930 | Alcohol<br>Hydroxy<br>Primary alcohol | No hits |
|  | Eicosanoic acid | 1647 | Straight chain fatty acid |  |  |  |
|  | Unkonwn 1669 | 1672 |  | 3023 | Alcohol<br>Hydroxy<br>Primary alcohol | No hits |
|  | Unknown Compound 1720 | 1721 |  | 3151 | Hydroxy<br>Alcohol<br>Carboxylic acid<br>Primary alcohol | splash10-05o0-1910000000-49e59c211689b393e2a6 |
|  | Unknown compound 1732 | 1734 |  | 3185 | Alcohol<br>Hydroxy<br>Primary alcohol | No hits |
|  | Unknown compound 1746 | 1746 |  | 3215 | Alcohol<br>Hydroxy<br>Primary alcohol | No hits |
|  | Unknown 2005 | 2005 |  | 3691 | Alcohol<br>Hydroxy<br>Alkene | No hits |
|  | Unknown 2030 | 2035 |  | NA | Hydroxy | No hits |

**Table S6.** Metabolite mixtures used across experiments. All mixtures contain a diverse range of compounds representing multiple metabolite classes. A reduced set of compounds were combined to both show the effects of salt and water on metabolite detection and create calibration curves for specific compounds.

| Compound | Class | Experiment Mixture |
| --- | --- | --- |
| Pyruvate | Alpha keto-acid | Method development and SPE |
| Alanine | Amino acid | Method development and SPE |
| Arginine | Amino acid | Method development and SPE |
| Aspartic acid | Amino acid | Method development and SPE |
| Carnitine | Amino acid | Method development and SPE |
| Cysteine | Amino acid | Method development and SPE |
| Glutamate | Amino acid | Method development and SPE |
| Glutamic acid | Amino acid | Method development and SPE |
| Glutamine | Amino acid | Method development and SPE |
| Glycine | Amino acid | Method development and SPE |
| Histidine | Amino acid | Method development and SPE |
| Isoleucine | Amino acid | Method development and SPE |
| Lactic acid | Amino acid | Method development and SPE |
| Leucine | Amino acid | Method development and SPE |
| Lysine | Amino acid | Method development and SPE |
| Methionine | Amino acid | Method development and SPE |
| Ornithine | Amino acid | Method development and SPE |
| Phenylalanine | Amino acid | Method development and SPE |
| Proline | Amino acid | Method development and SPE |
| Serine | Amino acid | Method development and SPE |
| Threonine | Amino acid | Method development and SPE |
| Thymine | Amino acid | Method development and SPE |
| Tryptophan | Amino acid | Method development and SPE |
| Valine | Amino acid | Method development and SPE |
| Citric acid | Organic acid | Method development and SPE |
| Fumerate | Organic acid | Method development and SPE |
| Lauric acid | Organic acid | Method development and SPE |
| Maleic acid | Organic acid | Method development and SPE |
| Malic acid | Organic acid | Method development and SPE |
| Succinic acid | Organic acid | Method development and SPE |
| DMSP | Organic compound | Method development and SPE |
| Oxalic Acid | Organic compound | Method development and SPE |
| Urea | Organic compound | Method development and SPE |
| Cellobiose | Sugar | Method development and SPE |
| Fructose | Sugar | Method development and SPE |
| Galactose | Sugar | Method development and SPE |
| Glucose | Sugar | Method development and SPE |
| Maltose | Sugar | Method development and SPE |
| Mannose | Sugar | Method development and SPE |
| NAG | Sugar | Method development and SPE |
| Ribose | Sugar | Method development and SPE |
| Sucrose | Sugar | Method development and SPE |
| Trehalose | Sugar | Method development and SPE |
| Mannitol | Sugar alcohol | Method development and SPE |
| Myo-inositol | Sugar alcohol | Method development and SPE |
| Glycerophosphate | Sugar phosphate | Method development and SPE |
| Alanine | Amino acid | Quantifying salt-water effect |
| Glutamine | Amino acid | Quantifying salt-water effect |
| Glycine | Amino acid | Quantifying salt-water effect |
| Lactate | Amino acid | Quantifying salt-water effect |
| Leucine | Amino acid | Quantifying salt-water effect |
| Lysine | Amino acid | Quantifying salt-water effect |
| Ornithine | Amino acid | Quantifying salt-water effect |
| Serine | Amino acid | Quantifying salt-water effect |
| Succinate | Amino acid | Quantifying salt-water effect |
| Thymine | Amino acid | Quantifying salt-water effect |
| Valine | Amino acid | Quantifying salt-water effect |
| Citric acid | Organic acid | Quantifying salt-water effect |
| Glutamic acid | Organic acid | Quantifying salt-water effect |
| Malic acid | Organic acid | Quantifying salt-water effect |
| Oxalic acid | Organic acid | Quantifying salt-water effect |
| Fructose | Sugar | Quantifying salt-water effect |
| Glucose | Sugar | Quantifying salt-water effect |
| NAG | Sugar | Quantifying salt-water effect |
| Sucrose | Sugar | Quantifying salt-water effect |
| Trehalose | Sugar | Quantifying salt-water effect |
| Myo-inositol | Sugar alcohol | Quantifying salt-water effect |
| Glycine | Amino acid | Quantify detection limits |
| Histidine | Amino acid | Quantify detection limits |
| Leucine | Amino acid | Quantify detection limits |
| Lysine | Amino acid | Quantify detection limits |
| Ornithine | Amino acid | Quantify detection limits |
| Serine | Amino acid | Quantify detection limits |
| Thymine | Amino acid | Quantify detection limits |
| Valine | Amino acid | Quantify detection limits |
| Citrate/citric acid | Organic acid | Quantify detection limits |
| Glutamic acid | Organic acid | Quantify detection limits |
| Lactic Acid | Organic acid | Quantify detection limits |
| Lauric acid | Organic acid | Quantify detection limits |
| Malic acid | Organic acid | Quantify detection limits |
| Succinate | Amino acid | Quantify detection limits |
| Cholesterol | Sterol | Quantify detection limits |
| Fructose | Sugar | Quantify detection limits |
| Glucose | Sugar | Quantify detection limits |
| Sucrose | Sugar | Quantify detection limits |
| Trehalose | Sugar | Quantify detection limits |
| Myo-inositol | Sugar alcohol | Quantify detection limits |

### Supplementary Text 1. R script for peak picking for GC-MS data

~~~
# Peak Picking.R
# EM Sogin
# Description: R script to pick peaks from GC-MS data

library(xcms)
library(CAMERA)

## PEAK PICKING, RETENTION TIME GROUPING & CORRECTION WITH XCMS
setwd(‘home/path/to/files’)
files<-list.files(pattern=‘.mzXML’, recursive = T, full.names=T)
xs <-xcmsSet(files, method = “matchedFilter”, fwhm = 8.4, snthresh = 1,step= 0.25, steps= 2,sigma =
3.56718192627824, max= 500, mzdiff= 1,index= FALSE)
xset1 <-group(xs,method = “density”, bw=2, mzwid= 1, minfrac = 0.3, minsamp = 1,max = 500) ##
Initial peak grouping
xset2 <-retcor(xset1)
xset2 <-group(xset2,method = “density”, bw=2, mzwid= 1, minfrac = 0.3, minsamp = 1,max = 500)
xset<-fillPeaks(xset2)

## Group peaks in to pseudo-spectra using CAMERA
an<-xsAnnotate(xset)
xsF<-groupFWHM(an, perfwhm=3)

peaks<-getPeaklist(xsF)
peaks[is.na(peaks)]<-0

save.image(‘Peak_Picking_Results.RData’)
# End
~~~

